# Genetic adaptation rates differ by trait and plant type --a comprehensive meta-analysis

**DOI:** 10.1101/2022.12.29.522194

**Authors:** Jianhong Zhou, Ellen Cieraad, Peter M. van Bodegom

**Affiliations:** Institute of Environmental Sciences (CML), Leiden University, 2333 CC Leiden, The Netherlands; Nelson Marlborough Institute of Technology, 322 Hardy Street, Nelson 7010, New Zealand

**Keywords:** evolutionary rate, genetic adaptation, global meta-analysis, intraspecific trait variation, plant characteristics, plant functional traits

## Abstract

- Plants can respond to changing climatic conditions through genetic adaptation of their functional traits. Despite the relevance of adaptation to climate change, much remains unknown about plant genetic adaptation rates, including whether these rates differ among plant characteristics and trait types.
- We performed a meta-analysis to investigate patterns of genetic adaptation rates and assess how these rates differ among plant species groups, growth forms and trait types, using a newly compiled database from 74 studies, comprising 35 functional traits across 72 angiosperm species. This database specifically focuses on genetic adaptation of plant functional traits, as separating from phenotypic plasticity.
- Both annual and generational adaptation rates of plant traits decrease non-linearly with increasing elapsed time since divergence. Plants adapt fastest when first introduced to a changing environment, but these rates go slow subsequently. Overall, shrubs have higher adaptation rates than trees, which confers shrubs an adaptive advantage over trees. Different adaptation rates among growth forms, life histories and trait types suggest an important additional mechanism through which climate change may affect community composition.
- Our study has important implications regarding plant adaptation to new environments and will improve the prediction of vegetation responses and ecosystem functioning upon climate change.

## 1. Introduction

Climate change is one of the major challenges of our time and will lead to profound impacts on biodiversity and ecosystem functions (IPCC, 2021). Plants, the fundamental part of ecosystems, play a vital role in maintaining biodiversity and ecosystem functioning. There has been a strong interest in how, and how fast, plants may respond to a changing environment (Hoffmann & Sgró, 2011; Anderson *et al*., 2013; Maire *et al*., 2013) and in particular whether certain plant species can adapt to future climatic conditions (Gienapp *et al*., 2008; Moran *et al*., 2016).

Plants can respond to changing climatic conditions through the variation of their functional traits, as functional traits are closely related to the performance of individual species, community assembly and ecosystem functioning (Webb *et al*., 2010; Laughlin *et al*., 2012; Maire *et al*., 2013; Van Bodegom *et al*., 2014). Trait variation includes the variation between species (interspecific trait variation) and within species (intraspecific trait variation, hereafter ITV). While previous studies mainly focused on interspecific trait variation (Wright *et al*., 2004; Reich, 2014; Díaz *et al*., 2016), recent studies (Li *et al*., 2018; Yang *et al*., 2020; Westerband *et al*., 2021) have shown that ITV also plays a critical role for plants in adapting to varying climatic conditions (Messier *et al*., 2010; Violle *et al*., 2012; Siefert *et al*., 2015; Malyshev *et al*., 2016; Henn *et al*., 2018; Zhou *et al*., 2022). Both phenotypic plasticity and genetic adaptation could drive ITV which is triggered by a changing climate (Abrams, 1994; Westerband *et al*., 2021). Phenotypic change through genetic adaptation involves the changing of genetic basis between generations while that through phenotypic plasticity does not (Abrams, 1994). Therefore, trait variation resulting from genetic adaptation could develop beyond the corresponding genotypic constraints of a species and thus the produced new generations may survive under more stressful abiotic conditions. Knowing how fast this process can occur and understanding how it relates to plant characteristics will enable us to predict potential evolutionary winners and losers under future climatic conditions (Hoffmann & Sgró, 2011). While phenotypic plasticity and genetic adaptation contribute to phenotypic change very differently, current estimates of phenotypic variation in literature (Westley, 2011; Alberti *et al*., 2017; Zhou *et al*., 2022) are commonly determined by the combination of both mechanisms, rather than separately. Consequently, the research focussing on genetic adaptation rates of plant functional traits is quite limited.

There are mainly two types of studies focusing on plant trait variations related to genetic adaptation. The first type comprises observational studies with a common garden experiment or reciprocal transplant experiment that attempts to isolate the trait variation caused by genetic adaptation (De Villemereuil *et al*., 2015). Most of these studies compare how much the shift in traits between the populations collected from the native and invasive ranges and focus on understanding how this trait variation helps these species establish invasive populations (e.g. Leger & Rice, 2003; Blair & Wolfe, 2004; Hierro *et al*., 2009; Shirk & Hamrick, 2014; van Boheemen *et al*., 2019; Pal *et al*., 2020). Although these studies can be used to assess genetic adaptation rates, they are usually focused on a single or a few species and fail to conclude a general pattern of which trait or species groups may adapt faster.

The other type comprises meta-analytical studies. Three meta-analytical studies have so far investigated the genetic adaptation rates of plant traits. Bone & Farres (2001) performed the first meta-analysis about plant genetic adaptation rates and observed that perennials adapted faster than annuals per generation and physiological traits adapted faster than morphological traits. Yet these conclusions were based on a relatively small database (including 78 observations of 26 species). Crispo *et al*. (2010) also assessed if evolutionary patterns differed among trait types across 20 species (including 13 plant species) and but their study mainly focused on whether the evolution of phenotypic change was adaptive. Gorné & Díaz (2019) synthesized a larger database of phenotypic change (containing 152 plant species) for meta-analysis. They observed a general pattern between the phenotypic change rate and elapsed time and concluded this pattern was independent of life span, growth form and trait type. But their study emphasised the evolutionary pathways through which phenotypic changes occur (Gorné & Díaz, 2019) and thus did not comprehensively present how the phenotypic change rates differ among those plant features. Furthermore, their database combined the phenotypic change caused by phenotypic plasticity and genetic adaptation and included observations that were not widely regarded as important functional traits (eg. number of leaves/flowers/seeds) or even not actual plant traits (eg. herbivore richness). To sum up, although these meta-analyses have more or less touched upon whether genetic adaptation rates differed among some plant characteristics (life span, growth form and trait type), their assertions might be premature as those meta-analyses were based on either small datasets (Bone & Farres, 2001; Crispo *et al*., 2010) or a database which does not strictly focus on genetic adaptation and functional traits (Gorné & Díaz, 2019). Moreover, so far there are no specific hypotheses about the relationships between the genetic adaptation rates and plant characteristics, thus a generic understanding of whether and how the genetic adaptation rates are related to plant features (e.g. growth form, life history, trait type, species group) is missing. Such knowledge is crucial when addressing how and how fast plant species may respond, and vegetation may shift, under future climatic conditions.

To fill these gaps, we first compiled a new plant genetic adaptation rate database including 72 angiosperm species and 35 functional traits from 74 studies. Our database specifically includes only plant trait variation driven by genetic adaptation and focuses on ecologically relevant traits. As far as we are aware, this database is the largest high-quality database, which, for the first time, allows the extraction of general trends about the genetic adaptation of plant functional traits. Then, we performed a meta-analysis to comprehensively investigate the general pattern between the genetic adaptation rates of plant functional traits and plant characteristics. In particular, we assessed how these genetic adaptation rates changed with time since introduction to a changing environment and differed across plant life history, growth forms, phylogenetic groups and trait types.

## 2. Materials and methods

### 2.1 Database preparation

We created our database in five steps. Firstly, we searched published literature in the Web of Science by using the following search strings 1 “(invasive OR exotic OR introduced) AND native AND (“common garden*” OR transplant) AND trait AND plant)” and search string 2 “(darwins AND plant AND trait)”, which resulted in 237 and 51 articles (by 2021-08-25), respectively. The first string was chosen to identify studies with common garden and reciprocal transplant experiments that analyse populations of the same plant species from different locations, and these almost exclusively relate to invasive species. The second string reflects studies that directly determined genetic adaptation rates (as expressed by ‘darwins’, explained below).

Secondly, we screened for articles that either had available genetic adaptation rates of plant traits or allowed us to calculate those genetic adaptation rates. This resulted in 237 articles from search string 1. From the results of search string 2, we only selected three review articles as the other 48 papers did not calculate the genetic adaptation rates of plant traits. We then applied a snowballing approach to these three selected review articles (Bone & Farres, 2001; Purugganan & Fuller, 2011; Gorné & Díaz, 2019) and found 64 original articles that were used to calculate their genetic adaptation rates. We then submitted all 301 articles to a rigorous quality control process to create a database that allows the estimation of genetic adaptation rates as precisely as possible. We only selected those data within each article that fulfilled eight data quality control criteria: (1) Plants had been grown in common garden or transplant experiments, or the study contained time series data of species trait values, to ensure that the trait variation was only caused by genetic adaptation; (2) For pairs of native and invasive populations, we selected only those data related to ancestral populations from the native range and the descendant populations in the invasive range; (3) If it was indicated that a species had multiple introduced histories, we separated the pairs of the native and invasive populations according to their introduction history; (4) If a species had different polyploids, we only selected the data of native and invasive species of the same polyploid to avoid including potential step changes of genetic adaptation due to polyploidy; (5) If multiple studies were about the same species and from the same authors, we checked whether the data of these studies were based on a same experiment and, if so, only selected one study to avoid adding repeated data; (6) We only selected functional traits which were related to ecological processes and widely used across the studies in our database; some traits such as height and biomass which are easily influenced by ontogeny were excluded; (7) In each study, we selected the functional traits for which measurements followed standard protocols; (8) For experiments with different treatments, we only used data from treatments that were the closest to natural conditions (e.g. in terms of temperature, nutrient and moisture availability and herbivory pressure). A full list of the final selection of 74 studies is shown in the Supporting Information (Table S1).

Thirdly, we compiled our database. We extracted a series of variables from the selected papers, including functional trait(s) measured, native and exotic locations of the species (as most of the species we collected were invasive species), trait values of species which had been collected in both the native and the exotic range, time since divergence, and adaptation rates (see our dataset for more details). Trait values were extracted directly from the publications, from available raw datasets as provided in the text, tables, appendices or associated online datasets. If trait values were only available in figures, we used WebPlotDigitizer (https://apps.automeris.io/wpd/) to extract the data. If the time since divergence was not exactly indicated in the original studies, we searched other literature or other online resources. For those studies that did not provide estimates of genetic adaptation rates, we estimated adaptation rates using the metric that is similar to darwins (Haldane, 1949; see details below). Lastly, we also updated species names to accepted names according to The Plant List (http://www.theplantlist.org/) using the R package *taxonstand* (Cayuela *et al*., 2012). This new plant trait genetic adaptation database combines data from around the world (Fig. 1) and contains 306 adaptation rate estimates for 72 angiosperm species and 35 functional traits.

**Fig. 1.**
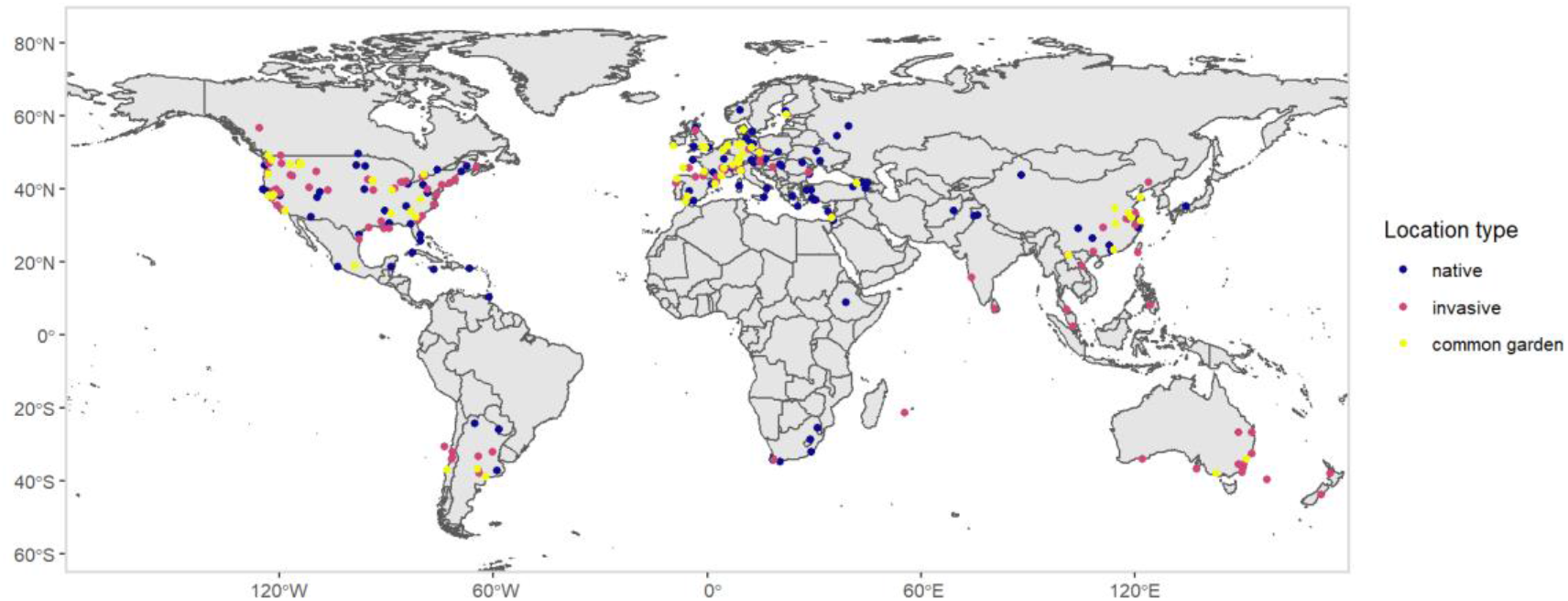
Native sample sites, invasive sample sites and locations of corresponding common garden experiments for 70 studies about invasive plant species in our global genetic adaptation database.

Fourthly, we added several more characteristics for each species: life span, generation time, life history, growth form and phylogenetic group, using a combination of existing datasets and online searches. For life span (annual, biennial or perennial), we first used the dataset “Plant life span” in the LEDA Traitbase (Kleyer *et al*., 2008, https://uol.de/en/landeco/research/leda/data-files), after updating species names of this dataset to accepted names using *Taxonstand*. For missing species, we completed these data using online searches on their life history information. Then, to allow correction for different numbers of generations per time period (and hence opportunities for genetic adaptation), we also estimated generation time, i.e. the amount of time (years) to reproductive age (Bone & Farres, 2001). For short-lived species, this was guided by their life span values: annuals have a generation time of 1 year and biennials have a generation time of 2 years. For species that were between annuals and biennials, we estimated their generation time as 1.5 years. For all perennial species, we performed online searches to source their generation time. For those species which mostly reproduce clonally, we did not estimate their generation time and recorded them as NA. Finally, we also categorized species into four life history categories based on their generation time: <1 year, 1-2 years, > 2 years and clonal.

To obtain growth form information for each species, we downloaded four public Categorical Plant Traits Databases from the TRY database (Kattge *et al*., 2011, dataset IDs: 57, 104, 110, 117; https://www.try-db.org/de/Datasets.php) and updated their species names to accepted names by using *Taxonstand*. Based on these four databases, we categorized species into four growth forms: herbs, graminoids (sedges, rushes and grasses), succulent (which was excluded in the statistical analysis as we only had one observation), shrubs and trees. If a growth form category for a species was inconsistent across the four databases or was not available, we performed an online search for information on the species.

To classify by phylogenetic groups, the family information of the species of our database (added using *Taxonstand*) was categorized according to the phylogenetic relatedness (close evolutionary nodes) indicated in the Angiosperm phylogeny (Cole *et al*., 2017). This resulted in seven phylogenetic groups: Poa-les (Poaceae), Cer-les (Ceratophyllaceae), Ran-les (Papaveraceae), Mal_Fab_Ros_Cuc-les (Euphorbiaceae, Hypericaceae, Phyllanthaceae, Fabaceae, Ulmaceae and Cucurbitaceae), Ger_Myr_Sap_Bra-les (Geraniaceae, Lythraceae, Sapindaceae and Brassicaceae), Car_Eri-les (Aizoaceae, Caryophyllaceae, Polygonaceae, Amaranthaceae, Ericaceae and Balsaminaceae) and Gen_Lam_Bor_Ast-les (Gentianaceae, Lamiaceae, Plantaginaceae, Boraginaceae and Asteraceae).

Lastly, we categorized the 35 functional traits into seven trait groups. The allocation trait group contained: root shoot ratio, reproductive allocation and vegetative shoot: aboveground biomass. The biochemistry group consisted of chlorophyll a content, total chlorophyll, leaf nitrogen content (LNC), leaf phosphorous content (LPC), leaf carbon content (LCC), leaf dry matter content (LDMC), leaf water content (LWC), leaf cellulose content, root nitrogen content, root phosphorous content and root carbon content. The ecophysiology group contained: CO_2_ assimilation rate per unit leaf area (A_area_), CO_2_ assimilation rate per unit leaf mass (A_mass_), maximum photosynthesis rate (P_max_), mean ambient photosynthetic rates (P_amb_), relative growth rate, growth rate, dark respiration rate (R_d_) and stomatal conductance (g_s_). The herbivory trait group was made up of the percentage of damaged leaf area by herbivore, percentage of damaged leave numbers per plant by herbivore, damaged leaf biomass by herbivore, seedling survival after herbivory and root regenerate survival after herbivory. The morphology group contained leaf area (LA), leaf mass per area (LMA), specific leaf area (SLA), leaf thickness, leaf density and root diameter. Fruit mass, seed mass and seed size were grouped as reproduction trait group. Gemination and survival comprised the survival trait group.

### 2.2 Estimating annual and generational adaptation rates

We estimated genetic adaptation rates of plant traits by the rate of evolutionary change suggested by Haldane (1949) below:

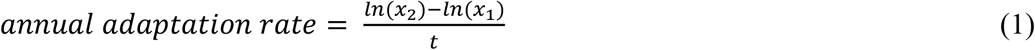

*x*_1_ is the mean trait value of the populations sampled in the native range (representing the ancestral populations), *x*_2_ is the mean trait value of the populations sampled in the introduced range (representing the descendant populations), *t* is the number of years elapsed since the ancestral population was introduced to a new environment until their descendant populations were collected in this new environment (if the collection time was not specifically indicated in their studies, we used the time when the common garden experiment was conducted instead). This annual evolutionary rate is equivalent to a millionth of a “darwins” as a unit of evolutionary rate.

For the two studies in our database with time series data (one collected from herbarium specimens (Buswell *et al*., 2011), the other from archaeological data of domestic plants (Purugganan & Fuller, 2011), we estimated, for each species, the average annual adaptation rates by plotting ln (*x*) values for each collected observation against the number of years (*t*) since the first collection, and then calculated the slope of a regression line through those observations as the adaptation rate (Hendry & Kinnison, 1999).

Since we expect genetic adaptation to primarily occur during reproduction, assessing the adaptation rates of species with different generation times may be problematic. We therefore also estimated generational adaptation rates by multiplying annual adaptation rates by generation time (Haldane, 1949). We propose that the generational adaptation rates are more closely related to the mechanism of genetic adaptation and represent the adaptability of plants more intrinsically.

### 2.3 Statistical analysis

Given that the direction of adaptation rates partly depends on how a trait is measured (e.g. LMA vs. SLA) and positive rates may not necessarily indicate improved fitness, we analysed the adaptation rates as absolute values. As the absolute annual and generational genetic adaptation rates (further referred to as annual and generational adaptation rates) were not normally distributed (Notes S1, Fig. S1, 2), we first used *descdist()* in *fitdistrplus* (Delignette-Muller & Dutang, 2015) package to find candidate distributions. Based on the skewness-kurtosis plot (Cullen & Frey, 1999; Fig. S1) provided by this function, we found these genetic adaptation rates more resembled γ, β or Weibull distribution. Then we used the *fitdist()* function in the same package to compare these distributions using Akaike Information Criterion (AIC). As the AIC value of Weibull distribution was the lowest, we concluded that the Weibull distribution fits the distribution of our annual and generational adaptation rates best.

We then performed a Generalized Linear Model (GLM) with Weibull distribution to test if the life history, growth form, phylogenetic groups and trait groups influenced annual and generational adaptation rates and used Tukey post-hoc tests to test which categories differed significantly from one another. Additional analyses (Notes S1) were performed to test the effect of common garden location (Fig. S3), field vs. greenhouse experimental condition (Fig. S4, 5), maternal effect (Fig. S6) and significant levels of compared trait variation (Fig S7, 8). The database was compiled in Microsoft Excel 2019 and all analyses were conducted in R (version 3.6.3, R Core Team (2020)).

**Fig. 2.**
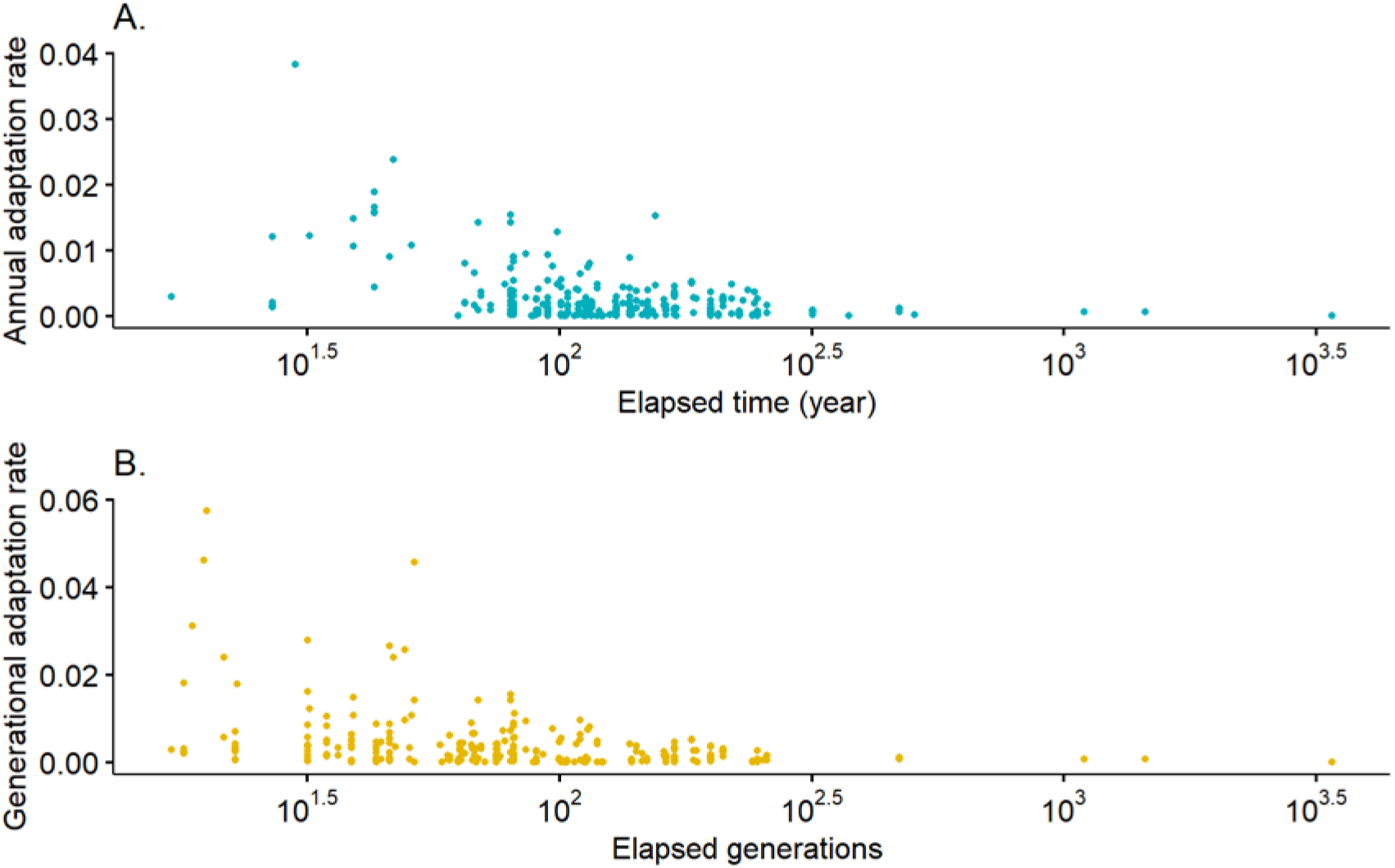
Annual adaptation rates of plant traits along time (A) and generational adaptation rates of plant traits along generations (B).

## 3 Results

Overall, annual adaptation rates of plant functional traits decreased with an increase in the time since divergence (Fig. 2A). The generational adaptation rates also decreased with the increasing number of generations (Fig. 2B).

Annual and generational adaptation rates were affected by plant life history categories (Fig. 3, R^2^ = 0.15 and R^2^ = 0.06, respectively). Regarding annual adaptation rates, clonal species had the highest rates, followed by early-matured, median-matured and later-matured plants, respectively. Late-matured plants had significantly lower annual adaptation rates than early-matured and clonal plants but did not differ from median-matured plants (Fig. 3A). In contrast, the generational adaptation rates of the late-matured plants were significantly higher than those of the early-matured plants, while both of their generational adaptation rates were not significantly different from those of median-matured plants (Fig. 3B).

**Fig. 3.**
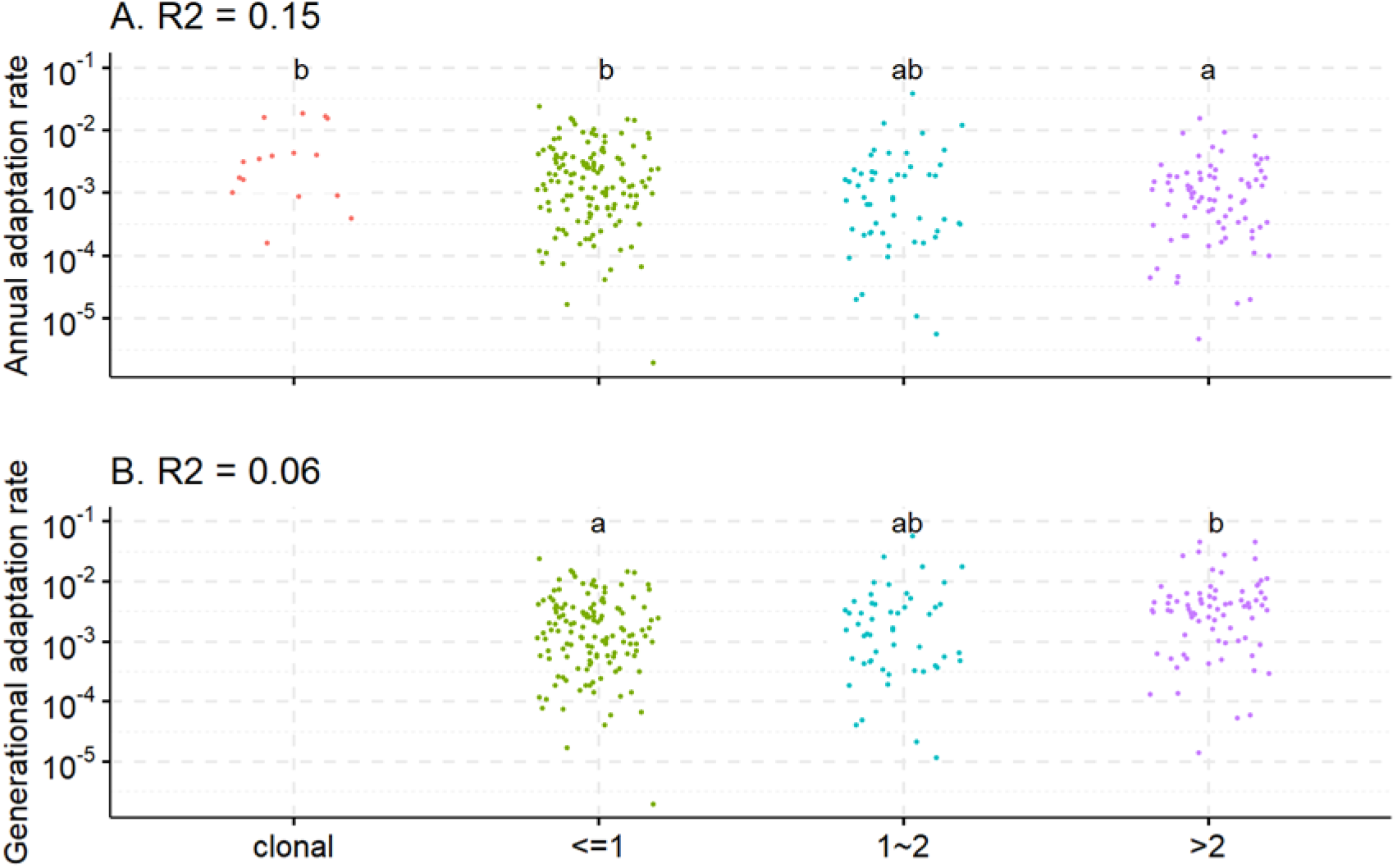
Patterns of annual (A) and generational (B) adaptation rates of plant traits among different life histories (clonal plants, generation time less than 1 year, generation time between 1 and 2 years, generation time more than 2 years). Different lowercase letters indicate that the means of plant annual or generational adaptation rates are significantly different between life history categories (as determined by Tukey post-hoc tests). When significant differences were detected, R^2^ indicates the explained variation, as estimated by Generalized Linear Model (GLM) with Weibull distribution. The generational adaptation rates of clonal plants are missing as their generation time are not available.

Annual adaptation rates were also different among different growth forms (Fig. 4A, R^2^ = 0.04). Shrub species had higher annual and generational adaptation rates than all other tested growth forms. Trees had significantly lower rates than herbs and shrubs while the rates of these three growth forms did not significantly differ from graminoids. By contrast, the generational adaptation rates were lowest for herbs (Fig. 4B, R^2^ = 0.08). Herbs had significantly lower adaptation rates than trees, shrubs and graminoids whereas there was no significant difference among the latter three growth forms.

**Fig. 4.**
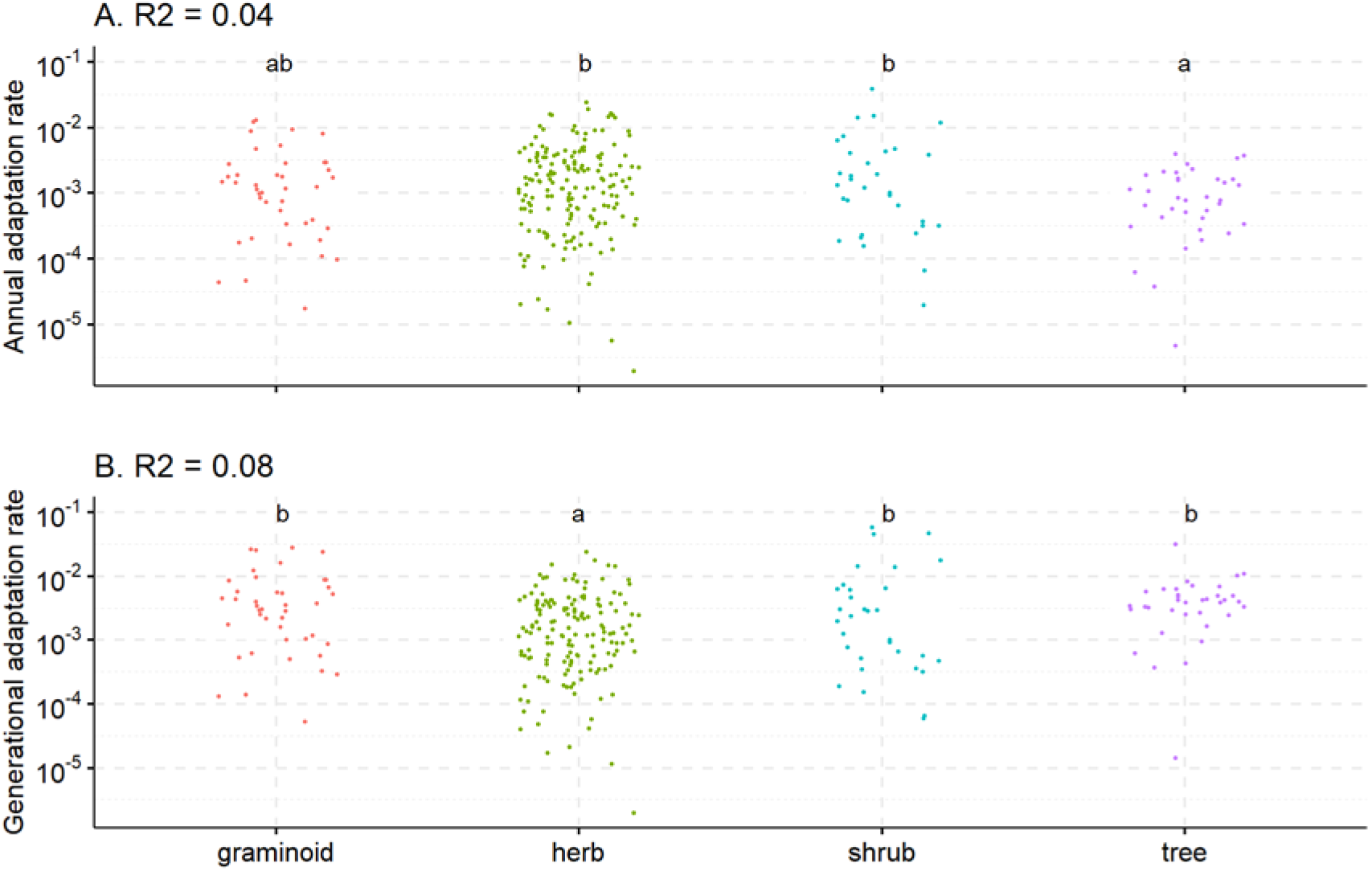
Patterns of annual (A) and generational (B) adaptation rates of plant traits among different growth forms. Different lowercase letters indicate that the means of plant annual or generational adaptation rates are significantly different between life history categories (as determined by Tukey post-hoc tests). When significant differences were detected, R^2^ indicates the explained variation, as estimated by Generalized Linear Model (GLM) with Weibull distribution.

Plant species also differed in their adaptation rates among phylogenetic groups (Fig. 5). The annual adaptation rates of Gen_Lam_Bor_Ast-les were significantly higher than those of Ger_Myr_Sap_Bra-les, Mal_Fab_Ros_Cuc-les and Run-les, but did not significantly differ from those of Car_Eri-les and Poa-les. There were no significant differences in annual adaptation rates among Car_Eri-les, Ger_Myr_Sap_Bra-les, Mal_Fab_Ros_Cuc-les and Run-les groups (Fig. 5A, R^2^ = 0.15). Although the annual adaptation rates of Cer-les tended to be higher than those of other groups (except for Gem_Lam_Bor_Ast-les), we lacked the statistical power to prove any significant difference because there were only five observations in Cer-les group. The generational adaptation rates of Poa-les were significantly higher than those of Ger_Myr_Sap_Bra-les while they were not significantly different from rates of Car_Eri-les, Gen_Lam_Bor_Ast-les and Mal_Fab_Ros_Cuc-les (Fig. 5B, R^2^ = 0.06). Similarly, the generational adaptation rates of Ran-les tended to be lower than those of Car_Eri-les, Gen_Lam_Bor_Ast-les, Mal_Fab_Ros_Cuc-les and Poa-les, we also lacked enough statistical power to prove this difference because there were only six observations in the Ran-les group.

**Fig. 5.**
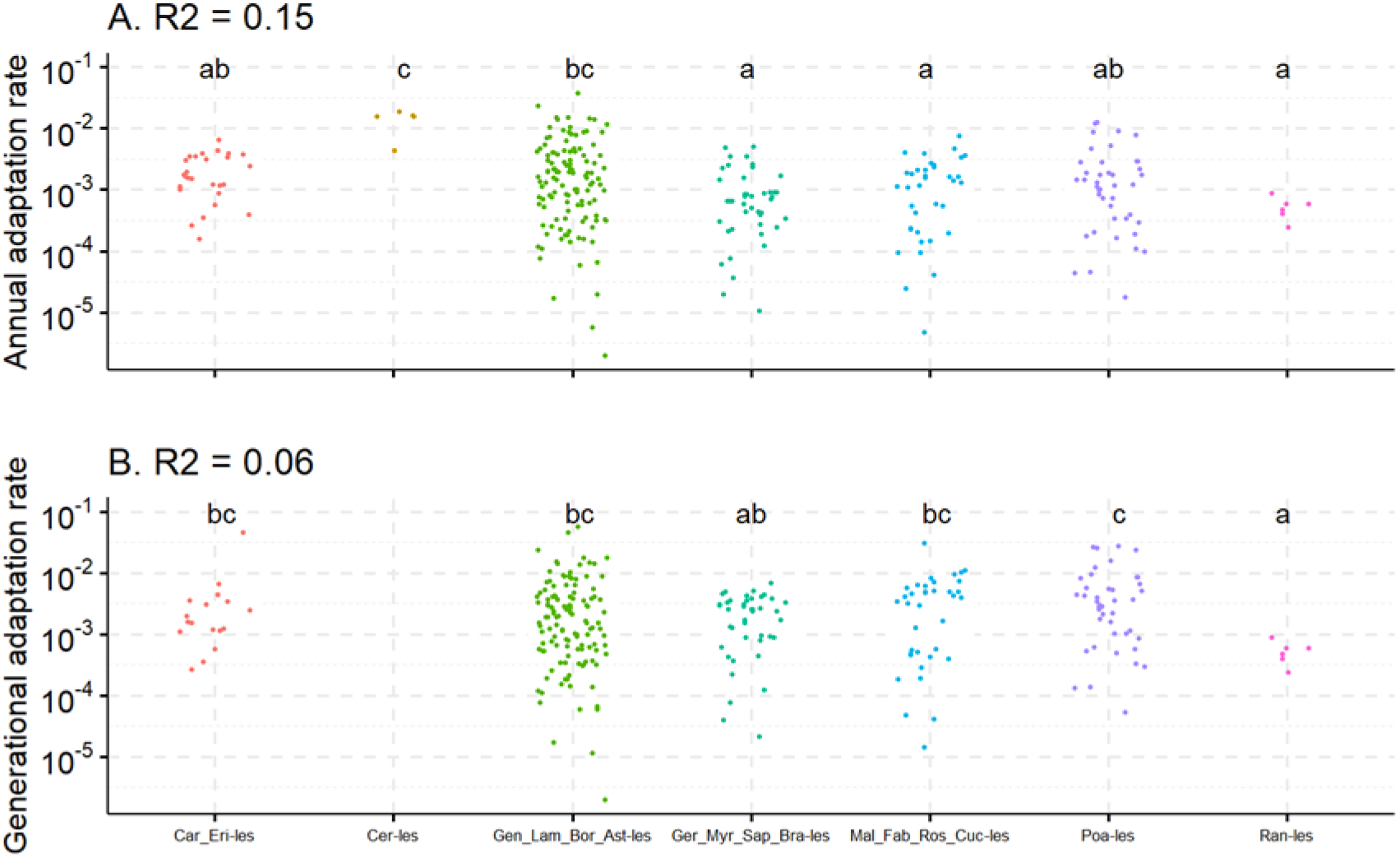
Patterns of annual (A) and generational (B) adaptation rates of plant traits among different phylogenetic groups. Different lowercase letters indicate that the means of plant annual or generational adaptation rates are significantly different between life history categories (as determined by Tukey post-hoc tests). When significant differences were detected, R^2^ indicates the explained variation, as estimated by Generalized Linear Model (GLM) with Weibull distribution. The generational adaptation rates of group Cer-les are missing as they are clonal plants and their generation time are not available.

Among different trait groups, biochemistry traits had the lowest annual adaptation rates (Fig. 6A, R^2^ = 0.07). Specifically, biochemistry adaptation rates were significantly lower than those of ecophysiology, allocation, survival and herbivory traits but not significantly lower than those of morphology and reproduction traits. These patterns remained similar for the mean generational adaptation rates among these trait groups (Fig. 6B, R^2^ = 0.13).

**Fig. 6.**
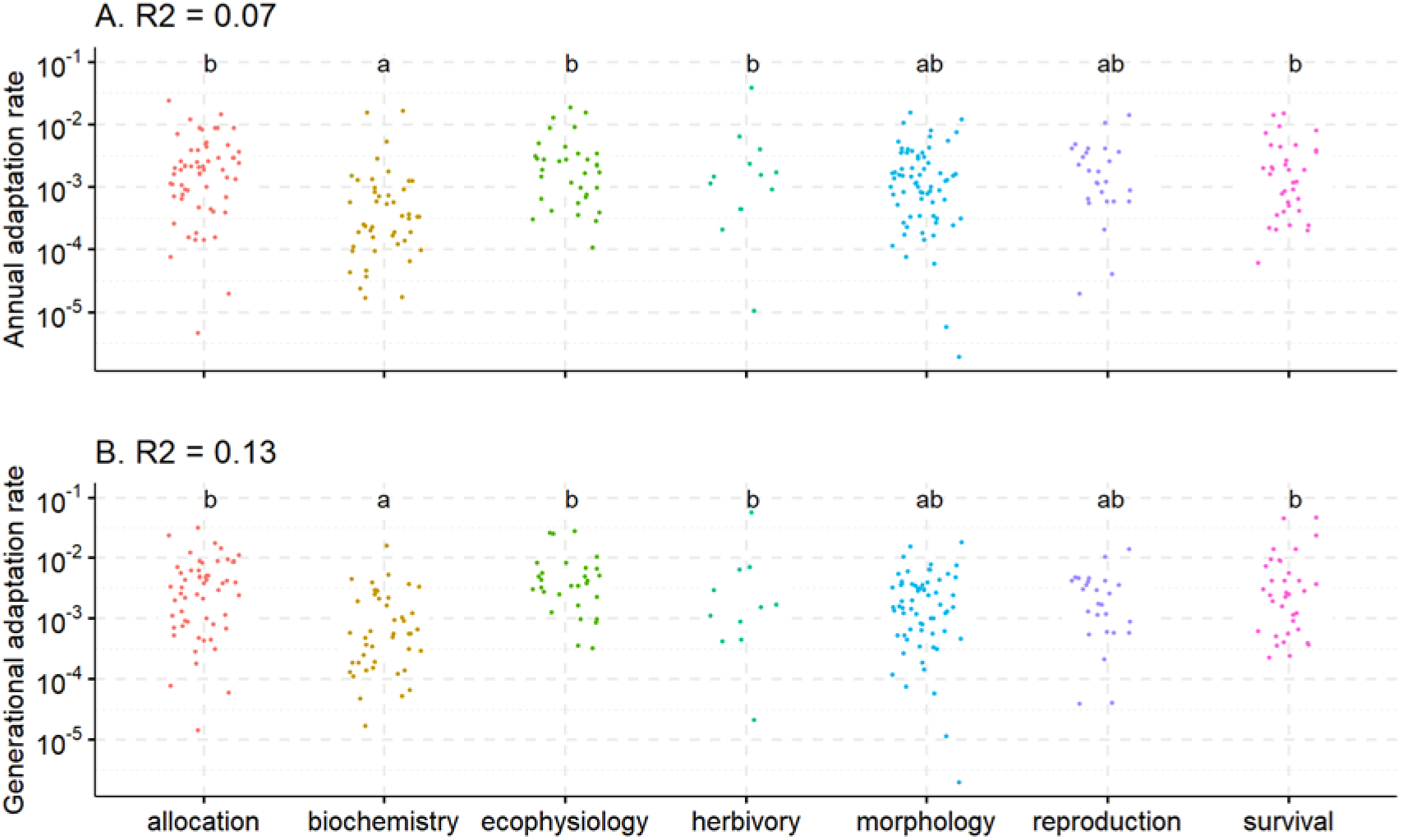
Patterns of annual (A) and generational (B) adaptation rates of plant traits among different trait groups. Different lowercase letters indicate that the means of plant annual or generational adaptation rates are significantly different between life history categories (as determined by Tukey post-hoc tests). When significant differences were detected, R^2^ indicates the explained variation, as estimated by Generalized Linear Model (GLM) with Weibull distribution.

## 4. Discussion

In this study, we evaluated both annual and generational adaptation rates as we consider these two metrics to have different implications for our understanding of genetic adaptation. For the annual adaptation rate, this is a metric which is widely used for calculating the adaptation rate (Hendry & Kinnison, 1999; Bone & Farres, 2001). On the one hand, it is a good measure for evaluating the adaptability of different plant species in the context of climate change, as they had to adapt to climate change within a given time period. On the other hand, the annual adaptation rate might not be a good reflection of the intrinsic differences in (mechanisms involved in) genetic adaptation rates of plants with different life history categories. As we expected that plants primarily adapt when they reproduced, we consider the adaptation rate per generation a more representative estimate of its mechanism. Our analyses indeed show the merits of making this distinction: while we showed that both annual and generational genetic adaptation rates were influenced by life history, growth form, phylogenetic group and trait type, the patterns between annual and generational adaptation rates were different. We observed that some patterns were even flipped between annual and generational adaptation rates.

### Patterns of adaptation rates

Assessing patterns in adaptation between plants with different life histories, we found that annual adaptation rates of earlier-matured plants were higher than those of later-matured plants, while generational adaptation rates of earlier-matured plants were lower than those of later-matured plants (Fig. 3). Not surprisingly, this suggests that annual and biennial plants have a higher annual potential to adapt to changing conditions simply because they reproduce earlier. On the other hand, the generational adaptation rates indicate that the intrinsic capacity for adaptation is higher for later-matured plants than for earlier-matured plants. This result complies with earlier findings that perennials have higher generational adaptation rates than annual plants (Bone & Farres, 2001). However, this intrinsic capacity is not able to compensate for the longer generation time, leading to overall lower annual adaptation rates. For clonal plants, their annual adaptation rates were as high as those of early-matured plants. While this may be considered strange for plants with a clonal strategy, it has been found that the three species with most observations in this category (*Alternanthera philoxeroides, Ceratophyllum demersum, Rorippa austriaca*) could occasionally propagate sexually (Dietz *et al*., 2002; Wang *et al*., 2005; Syed *et al*., 2018). If true, then their life history categories resemble those of early-matured species, which might explain their similarity in adaptation rates.

Comparing annual and generational adaptation rates between growth forms, we also found reversed patterns. Overall, trees had lower annual adaptation rates but higher generational adaptation rates than those of herbs (Fig. 4). To some extent, these results are consistent with the life history results, as trees usually mature later than herbs (in our database, herbs matured within two years while trees matured after more than two years, Fig. S9). These results imply that, although trees adapt faster per generation than herbs, within the same period of time, herbs are still adapting more quickly by producing more generations per time period than trees. Interestingly, life history does not fully compensate for their pattern of generational adaptation rates, as shrubs had the highest annual and generational adaptation rates among these growth forms (Fig. 4). This suggests that shrubs have a high capacity to adapt both at annual and generational levels. When looking in more detail at these results by splitting the patterns for different life history categories (Fig. S9), the higher adaptation rates of shrubs were more obvious with an increase in generation time. That is, for median-matured plants, the annual adaptation rates of shrubs were significantly higher than herbs. For late-matured plants, both annual and generational adaptation rates of shrubs were significantly higher than trees (although there were only seven late-matured shrub observations in our database). This finding also supports previous modelling hypotheses which indicate that, compared with trees, the shrub growth form is more adaptive (Götmark *et al*., 2016).

Evaluating annual and generational adaptation rates between phylogenetic groups, we observed that annual adaptation rates of Gen_Lam_Bor_Ast-les were higher than those of all the other groups (except for Ber-les which only had five observations) while the generational adaptation rates of Gen_Lam_Bor_Ast-les became lower than those of Poa-les (Fig. 5). This coincides with the observation above that early-matured plants had higher annual adaptation rates while late-matured plants had higher generational adaptation rates. As the detailed patterns for different life histories indicated that the majority of observations from the Gen_Lam_Bor_Ast-les group were from early-matured species, while those in the Poa-les group were mainly from late-matured species (Fig. S10). However, the plant history did not explain all of the patterns. we noticed that the annual adaptation rates of Ran-les were significantly lower than those of Gen_Lam_Bor_Ast-les within the early-matured plants and the annual adaptation rates of Ger_Myr_Sap_Bra-les were significantly lower than those of the other phylogenetic groups (except for Car_Eri-les but there was only one observation in this group) within late-matured plants. Currently, we do not fully understand why Ran-les and Ger_Myr_Sap_Bra-les had lower annual adaptation rates, but may partially be driven by the coarse plant life history classes used (less than 1 year, between 1 and 2 years, more than 2 years). More representative observations across life history categories and phylogenetic groups would allow a more thorough investigation to understand these different phylogenetic patterns.

For different trait groups, we found that, generally, the annual and generational adaptation rates of biochemistry traits were the lowest and significantly different from all the other traits except for morphology and reproduction traits (Fig. 6). This pattern was less influenced by the plant life history. With increasing generation time, fewer trait groups had significantly higher annual adaptation rates than biochemistry traits, but adaptation rates of ecophysiology traits were always significantly higher than those of biochemistry traits (Fig. S11). Biochemistry traits had the lowest genetic adaptation rates might because this trait group may be more susceptible to environmental conditions and could be adapted easily by plasticity, which would allow the reallocation of energy for genetically adapting other traits that are more important for plant fitness and survival.

### General implications

Our study has important implications regarding plant adaptation to new environments emerging from climate change. Generally, we found that both annual and generational adaptation rates decreased with an increase in the time since divergence (Fig. 2). This indicates that plants may adapt much faster when first faced with a different or changing environment, compared with after they have established adapted populations in the new environment. This finding is consistent with previous modelling result which indicated that plant adaptation happened instantaneously, following a change to a novel environment and then stayed stable (Gorné & Díaz, 2019). This may also partly explain why the observed plant speciation rates during the Anthropocene were much higher than the background rates (Thomas, 2015) and implies that plants may adapt, at least initially, to climate change rather quickly. These substantially increased adaptation rates above background levels also suggest that the estimated biodiversity loss caused by climate change might have been overestimated in some scenario studies (Thuiller *et al*., 2005).

We also showed that the adaptation rates were not constant within the plant kingdom as the adaptation rates differed depending on the life history, growth form, phylogeny and trait group. Given that local adaptation is considered critical to survival upon climate change (e.g. Aitken *et al*., 2008; Hoffmann & Sgró, 2011), these differences in adaptation rates also suggest that some species are more likely to avoid extinction – thanks to local adaptation – than others. Apart from the question of whether the new habitat conditions are suitable for a given species, determining the demand for local adaptation is likely to cause winners and losers of climate change. Our results show that trees have the lowest annual adaptation rates while those of shrubs were the highest. Shrubs may therefore have an advantage over trees in adapting to climate change. This advantage and disadvantage among growth forms may lead to major shifts in plant community composition in the future.

Our study is the first to directly assess differences in adaptation rates among different trait groups. This may be particularly important since future climatic drivers may act as environmental filters to select certain trait combinations, but this process could be constrained if there are uneven adaptation rates. For example, elevated atmospheric CO_2_ concentrations combined with warmer temperatures may select for ecophysiological traits that enable faster growth to counteract higher respiration demands induced by increased temperatures and more frequent drought may select higher root: shoot ratios to reduce water loss (Tjoelker *et al*., 1999; Karcher *et al*., 2008; Chen *et al*., 2022). We found that most trait groups had similar adaptation rates. The exception were biochemistry traits – these had lower adaptation rates but this may be compensated for by higher plasticity rates for this group. Overall, the lack of differences between trait groups suggests that the adaptation towards demanded trait combinations will unlikely be constrained by uneven trait adaptation rates.

### Conclusion

Our study, to our knowledge, is the first to comprehensively assess the general patterns between genetic adaptation rates and plant characteristics. The results have important implications regarding plant adaptation to new environments emerging from climate change. Higher adaptation rates following introduction to a changing environment suggests that plants may be able to adapt quite fast to changing climatic conditions. Differences in annual adaptation rates between growth forms imply that global change may lead to shifts in community composition. If our findings are generally true for extended datasets that similarly specifically target genetic adaptation rates (as distinct from phenotypic plasticity), these patterns will improve model prediction of vegetation responses and ecosystem functioning to climate change.

## Data availability

The data that supports the findings of this study will be openly available in the Zenodo repository upon acceptance.

## Acknowledgements

J Zhou was funded by a PhD scholarship from China Scholarship Council (No. 201608310102). We thank Lotte A. van Boheemen, José L. Hierro, Rebecca Y. Kartzinel for additional detailed information when we compiled some of their data into our database. We thank Xiaoyang Zhong for his insights to improve this manuscript.

## Author contributions

P.M.v.B conceived the study; J.Z., E.C. and P.M.v.B developed the ideas; J.Z. collected the data; J.Z. and E.C. conducted the analysis; J.Z. wrote the first draft. All authors contributed critically to the drafts and gave final approval for publication.

## Supporting Information

**Table S1** All studies from which data were compiled in our database

**Notes S1** Additional methodological details for data and statistical quality assurance

**Fig. S1** Skewness-kurtosis plot of annual (left) and generational (right) adaptation rates of plant traits (boot 10.000 times)

**Fig. S2** Frequency distribution of annual (A) and generational (B) adaptation rates of plant traits

**Fig. S3** Annual (A) and generational (B) adaptation rates of plant traits from the common gardens located in different locations

**Fig. S4** Annual (A) and generational (B) adaptation rates of plant traits based on different experimental conditions

**Fig. S5** Annual (A, B, C) and generational (D, E, F) adaptation rates of plant traits based on greenhouse or field experiments split by life history categories

**Fig. S6** Influence of maternal effect on annual (A) and generational (B) adaptation rates of plant traits

**Fig. S7** Annual (A) and generational (B) adaptation rates of plant traits with different significant levels in the trait variation between the native and invasive plants from the original studies

**Fig. S8** Annual (A, B, C) and generational (D, E, F) adaptation rates of plant traits with different significant levels in the trait variation between the native and invasive plants from their original studies split by life history categories

**Fig. S9** Patterns of annual (A, B, C) and generational (D, E, F) adaptation rates of plant traits among different growth forms split by life history categories

**Fig. S10** Patterns of annual (A, B, C) and generational (D, E, F) adaptation rates of plant traits among different phylogenetic groups split by life history categories

**Fig. S11** Patterns of annual (A, B, C) and generational (D, E, F) adaptation rates of plant traits among different trait groups split by life history categories

