## Supplemental Information for "Genetic adaptation rates differ by trait and plant type --a comprehensive meta-analysis"

The following Supporting Information is available for this article:

**Table S1 All studies from which data were compiled in our database**

| **study** | **Data source** | **reference** |
| --- | --- | --- |
| 1 | Table 2 | Lamarque *et al.* (2014) |
| 2 | Table 2 | Bone & Farres (2001) |
| 3 | Extracted from Fig. 4 | Hirsch *et al.* (2016) |
| 4 | Table 2 | Sternberg (2016) |
| 5 | published raw data | Ramula & Kalske (2020) |
| 6 | Published raw data | Li *et al.* (2020) |
| 7 | Published raw data | Sun & Roderick (2019) |
| 8 | Published raw data | Brandenburger *et al.* (2019) |
| 9 | Table 2, 3 | Pyšek *et al.* (2019) |
| 10 | Tables 2, 3, 4, 5 | Tho *et al.*, (2016) |
| 11 | Table 2 | Guo *et al.*, (2016) |
| 12 | Extracted from Fig. 2c (control) | Kleine *et al.* (2017) |
| 13 | Extracted from Fig. 1b, c &2a | Rouifed *et al.* (2018) |
| 14 | Extracted from Fig. 3a, c | Hernández *et al.* (2019) |
| 15 | Extracted from Fig. 2a, b, Fig. 3a, Fig. 4b, c, d | Gruntman *et al.* (2020) |
| 16 | Extracted from Fig. S2 | Pal *et al.* (2020) |
| 17 | Extracted from Fig. 2e, d | Liao *et al.* (2014) |
| 18 | Extracted from Fig. 2a, b | Zheng *et al.* (2013) |
| 19 | Extracted from Fig. 3a, Fig. 5a | Liao *et al.* (2013) |
| 20 | Published raw data | Turner *et al.* (2015) |
| 21 | published raw data | Zhang *et al.* (2015) |
| 22 | Published raw data | Shirk & Hamrick (2014) |
| 23 | Extracted from Fig. 1a, b | Oduor *et al.* (2013) |
| 24 | Extracted from Fig. S1 | Shang *et al.* (2015) |
| 25 | Extracted from Fig. 2, 3, 4, 5 | Lázaro-Lobo *et al.* (2020) |
| 26 | Table A3 | Hierro *et al.* (2013) |
| 27 | Extracted from Fig. 1B, F, I | Gard *et al.* (2013) |
| 28 | Extracted from Fig. 1a, b, e | Hyldgaard & Brix (2012) |
| 29 | Table 1 | Guo *et al.* (2011) |
| 30 | Extracted from Fig. 1 d, g | Chun (2011) |
| 31 | Extracted from Fig. 4c, i, q, Fig. 6a | Buswell *et al.* (2011) |
| 32 | Extracted from Fig. 1c, Fig. 2b, e | Oduor *et al.* (2011) |
| 33 | Extracted from Fig. 2A, Fig. 5B | Molina-Montenegro *et al.* (2011) |
| 34 | Extracted from Fig. 3e, g | Huang *et al.* (2010) |
| 35 | Extracted from Fig. 4 | Erfmeier & Bruelheide (2010) |
| 36 | Table A1 | Hierro *et al.* (2009) |
| 37 | Extracted from Fig. 1A, C | Quiroz *et al.* (2009) |
| 38 | Extracted from Fig. 1b | Zou *et al.* (2008) |
| 39 | Table 2 | Zou *et al.* (2007) |
| 40 | Extracted from Fig. 1, 2, 3 | Maron *et al.* (2007) |
| 41 | Extracted from Fig. 3f, i, j | Leger & Rice (2007) |
| 42 | Extracted from Fig. 1 | Leger & Rice (2003) |
| 43 | Table 1 | Güsewell *et al.* (2006) |
| 44 | Extracted from Fig. 2; Table 1 | Blair & Wolfe (2004) |
| 45 | Published raw data | Colomer-Ventura *et al.* (2015) |
| 46 | Extracted from Fig. 4, Fig. 5a | Hodgins & Rieseberg (2011) |
| 47 | Published raw data | Yang *et al.* (2021) |
| 48 | Published raw data | Van Boheemen *et al.* (2019) |
| 49 | Published raw data | Tewes & Müller (2018) |
| 50 | Published raw data | Gruntman *et al.* (2017) |
| 51 | Extracted from Fig. 1c, 3b | Atlan *et al.* (2015) |
| 52 | Extracted from Fig. 2a, b | Henery *et al.* (2010) |
| 53 | Extracted from Table 1 | Schlaepfer *et al.* (2010) |
| 54 | Extracted from Fig. 1 | Monty *et al.* (2010) |
| 55 | Extracted from Fig. 1 | Young *et al.* (2007) |
| 56 | Extracted from Fig. 3 | Roiloa *et al.* (2016) |
| 57 | Extracted from Fig. S2 | Meimberg *et al.* (2010) |
| 58 | GRAIN/SEED SIZE DATA in supplementary material | Purugganan & Fuller (2011) |
| 59 | Published raw data | Gorné & Díaz (2019) |
| 60 | Extracted from Fig. 2c, Fig. 3a, b | Seifert *et al.* (2009) |
| 61 | Table 3 | Allan & Pannell (2009) |
| 62 | Table 3 | Bossdorf *et al.* (2004) |
| 63 | Extracted from Fig. 2, 3, 4a | Buschmann *et al.* (2005) |
| 64 | Extracted from Fig. 1 | Mu (2005) |
| 65 | Table 3 | Willis *et al.* (2000) |
| 66 | The first paragraph of the results in the main text | Wolfe *et al.* (2004) |
| 67 | Extracted from Fig. 5, 6 (red dots) | Vandegrift *et al.* (2015) |
| 68 | Table S5 | Turner *et al.* (2014) |
| 69 | Extracted from Fig. 1b, Fig. 2c | García *et al.* (2013) |
| 70 | Extracted from Fig. 1A, C, E, F, Fig. 2C, E | Moroney *et al.* (2013) |
| 71 | Extracted from Fig. 1 | McKenney *et al.* (2007) |
| 72 | Extracted from Fig. 1 | Willis *et al.* (1999) |
| 73 | Table 2 | Caño *et al.* (2009) |
| 74 | Table 4 | Daehler & Strong (1997) |

**Methods S1 Data and statistical quality control procedures (including Note S1, Fig. S1-S8)**

**Notes S1** **Additional methodological details for data and statistical quality assurance**

As the absolute annual and generational genetic adaptation rates (further referred to as annual and generational adaptation rates) were not normally distributed (Notes S1, Fig. S1), we first used *descdist*() in the *fitdistrplus* package (Delignette-Muller &Dutang, 2015) to find candidate distributions. Based on the skewness-kurtosis plot (Cullen &Frey, 1999) provided by this function, we found these genetic adaptation rates more resembled γ, β or Weibull. Then we used the *fitdist()* function in the same package to compare these distributions using Akaike Information Criterion (AIC). For both response variables, the AIC value of Weibull distribution was the lowest, hence we concluded that the Weibull distribution fits the distribution of our annual and generational adaptation rates best.

We expected that some of the conditions related to the study design (e. g. whether the location of the common garden was in a native or invasive range, the common garden experiment was conducted in a greenhouse or the field, some studies used F1 generations of sampled seeds for the common to eliminate maternal effect but others directly used the parent sampled seeds) may affect the results. Hence, we checked if the annual and generational adaptation rates were influenced by these additional factors.

- The location of the common garden did not significantly influence the annual and adaptation rates of plant functional traits (Fig. S3).
- The type of common garden experiment (i.e. a greenhouse experiment, or an outdoor or field experiment) slightly (R^2^=0.06) influenced the annual adaptation rates of plant functional traits (Fig. S4).
- Given the potential bias in the types of plants tested in the different settings, we assessed whether the influence of the type of experiment was affected by the plant life history (Fig. S5). Indeed, annual adaptation rates of late-matured plants grown in greenhouse experiments were higher than the plants growing in the field or outdoor (Fig. S5C).
- The type of seeds used (i.e. sampled parental seeds or new F1 seeds) did not significantly influence the annual and generational adaptation rates of plant functional traits (Fig. S6).

In addition, considering authors or journals may tend to publish the studies with significant statistical differences found, we also checked if the data we selected involved this publication bias. For those studies that presented significance values (whether significant or not) for the relevant native *vs*. invasive comparisons, we found that those with significant comparisons had higher annual and generational adaptation rates than those of non-significant comparisons (Fig. S7). This result was to be expected as a higher trait value difference would be found if the trait variation was significant. Those native vs invasive comparisons for which significance levels were not available, also had higher annual adaptation rates than non-significant comparisons. This was also reasonable as these unknown comparisons are likely to include both significant and non-significant comparisons. When looking in more detail at these results by splitting the patterns across different life history categories (Fig. S8), the patterns were similar to the patterns shown in Fig. S7. The comparisons that comprise the database were fairly evenly allocated between the three categories (significant, non-significant and no significance data available). Together, this suggests that our dataset is not strongly influenced by publication bias.

**Fig. S1** **Skewness-kurtosis plot of annual (left) and generational (right) adaptation rates of plant traits (boot 10.000 times)**

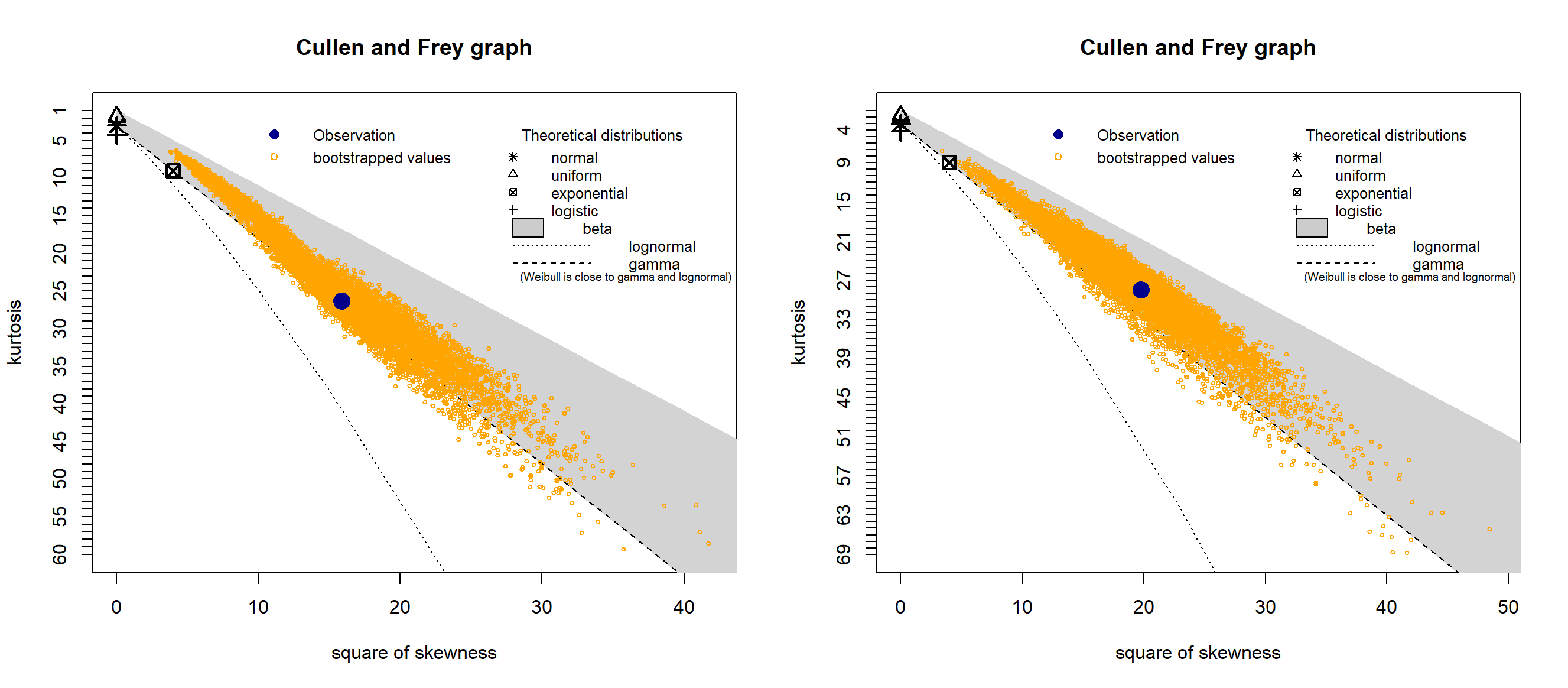

**Fig. S2** **Frequency distribution of annual (A) and generational (B) adaptation rates of plant traits**

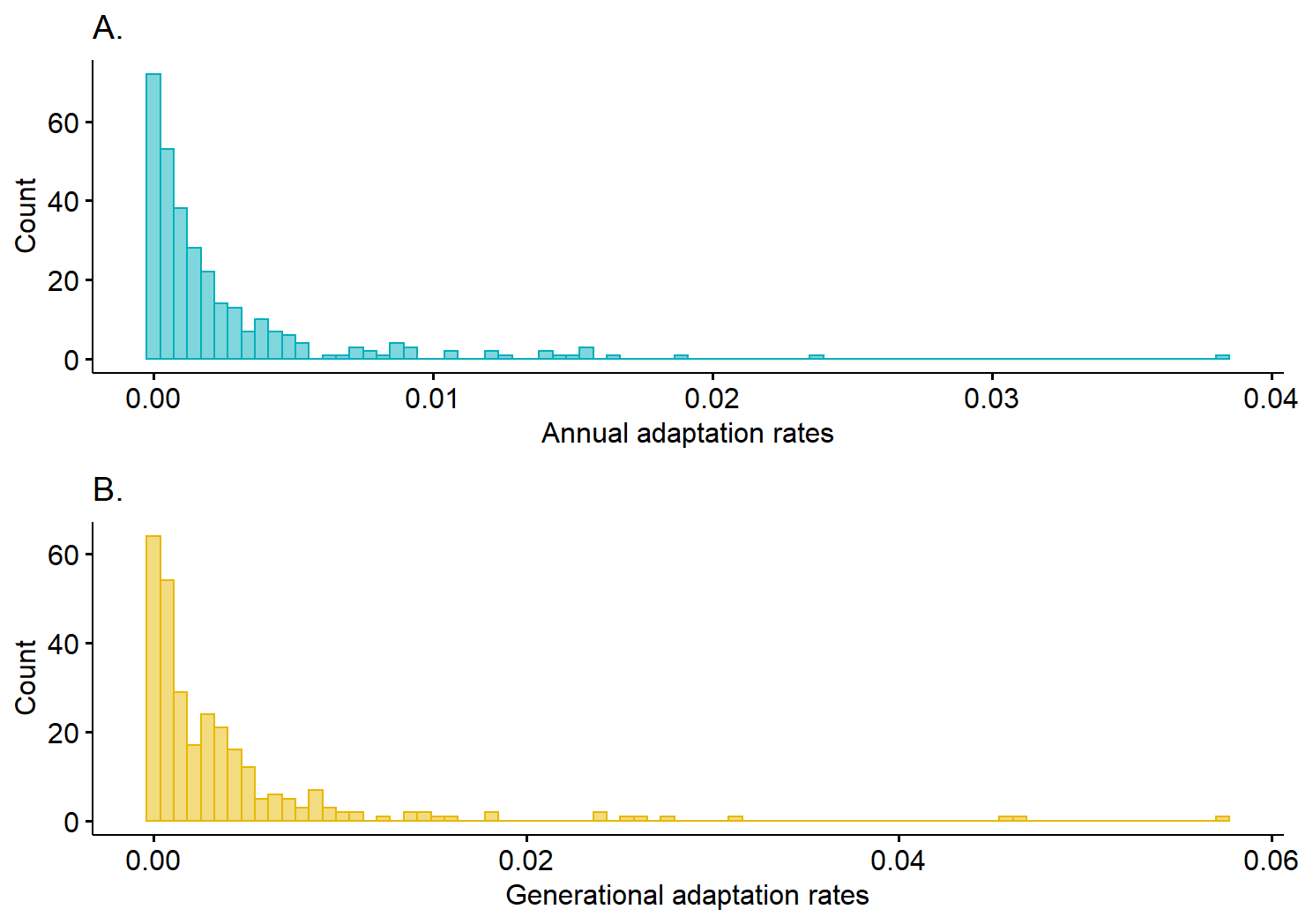

**Fig. S3 Annual (A) and generational (B) adaptation rates of plant traits from the common gardens located in different locations:** native country, invasive country, or third country (i.e. neither native or invasive country). Different lowercase letters indicate that the means of plant annual or generational adaptation rates are significantly different between life history categories (as determined by Tukey post-hoc tests).

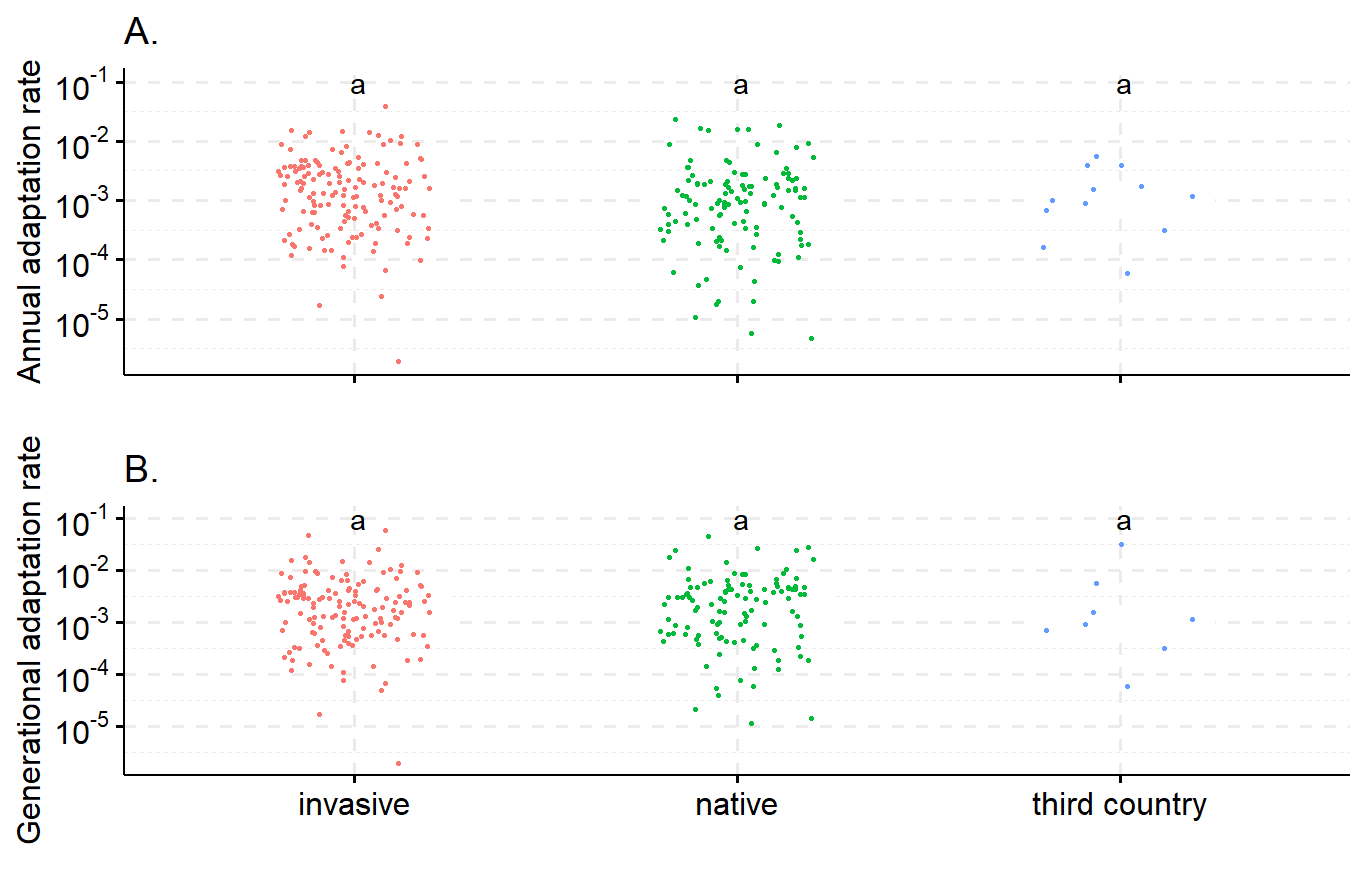

**Fig. S4 Annual (A) and generational (B) adaptation rates of plant traits based on different experimental conditions** (greenhouse experiment, field or outdoor experiments). Different lowercase letters indicate that the means of plant annual or generational adaptation rates are significantly different between life history categories (as determined by Tukey post-hoc tests). When significant differences were detected, R^2^ indicates the explained variation, as estimated by Generalized Linear Model (GLM) with Weibull distribution.

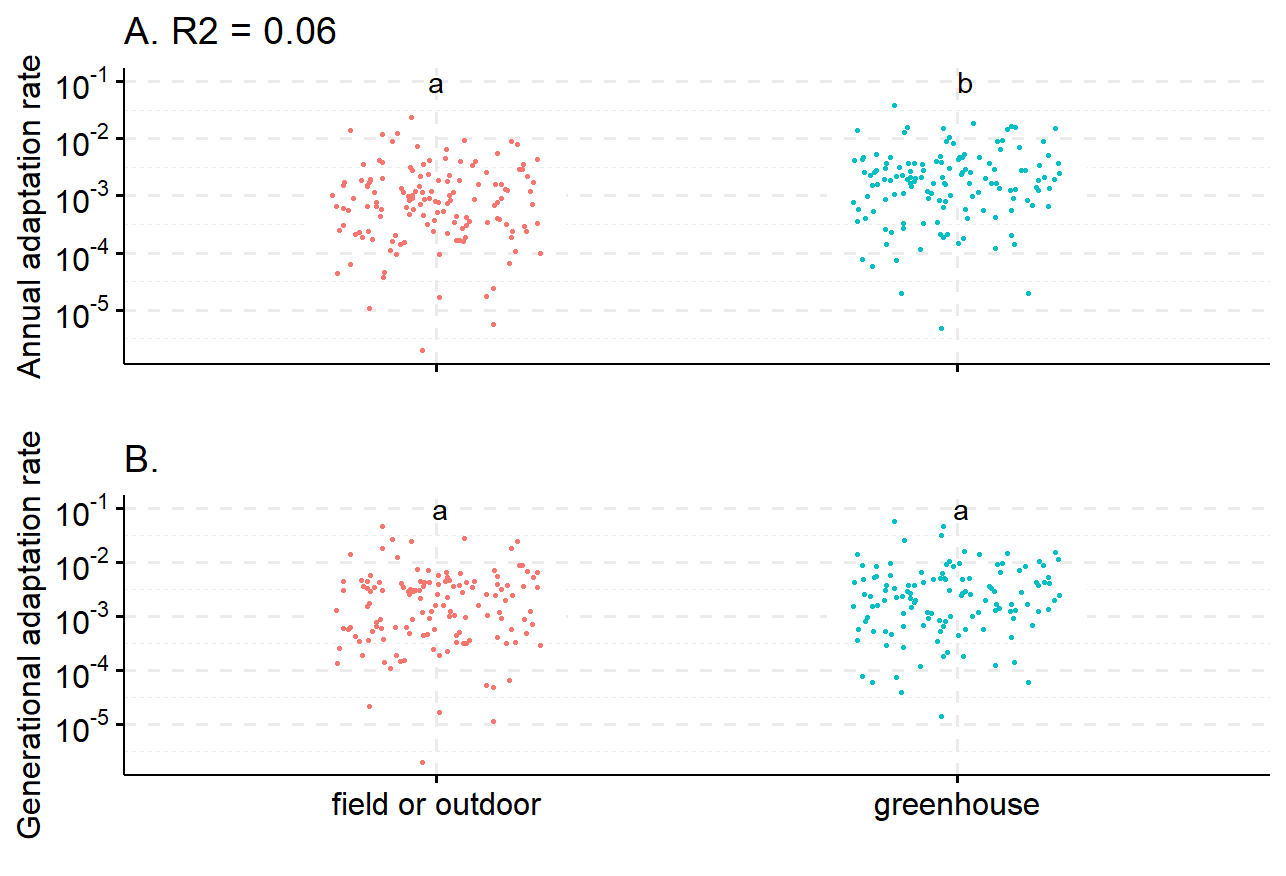

**Fig. S5 Annual (A, B, C) and generational (D, E, F) adaptation rates of plant traits based on greenhouse or field experiments split by life history categories** (A & D. generation time less than 1 year, B & E. generation time between 1 and 2 years, C & F. generation time more than 2 years). Different lowercase letters indicate that the means of plant annual or generational adaptation rates are significantly different between life history categories (as determined by Tukey post-hoc tests). When significant differences were detected, R^2^ indicates the explained variation, as estimated by Generalized Linear Model (GLM) with Weibull distribution.

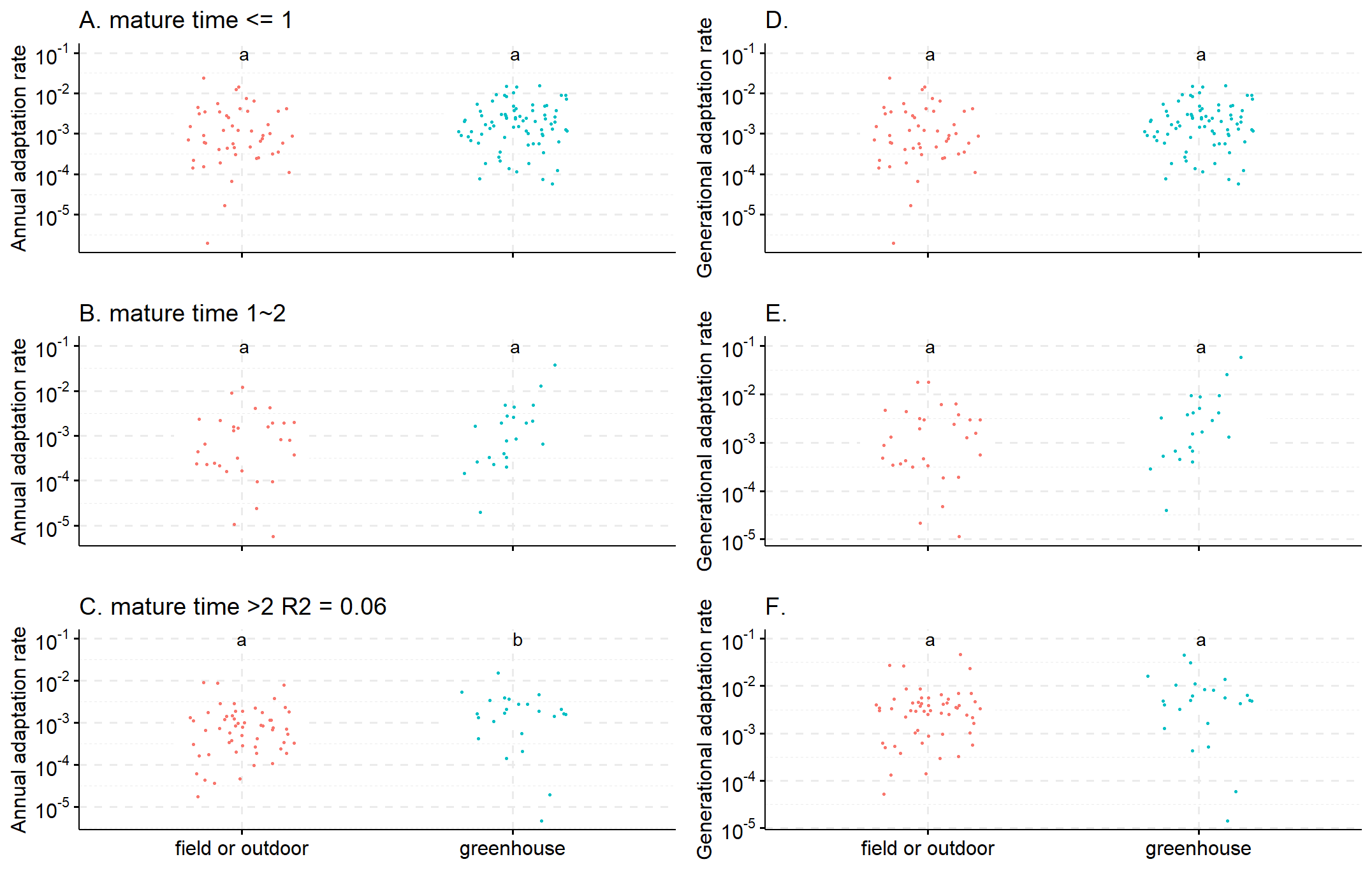

**Fig. S6 Influence of maternal effect on annual (A) and generational (B) adaptation rates of plant traits**. **‘**Yes’, indicates studies where the maternal effect was eliminated by using the next generation (F1) of seeds. ‘No’, maternal effect was not eliminated as sampled seeds were directly used in the common garden experiments. Different lowercase letters indicate that the means of plant annual or generational adaptation rates are significantly different between life history categories (as determined by Tukey post-hoc tests).

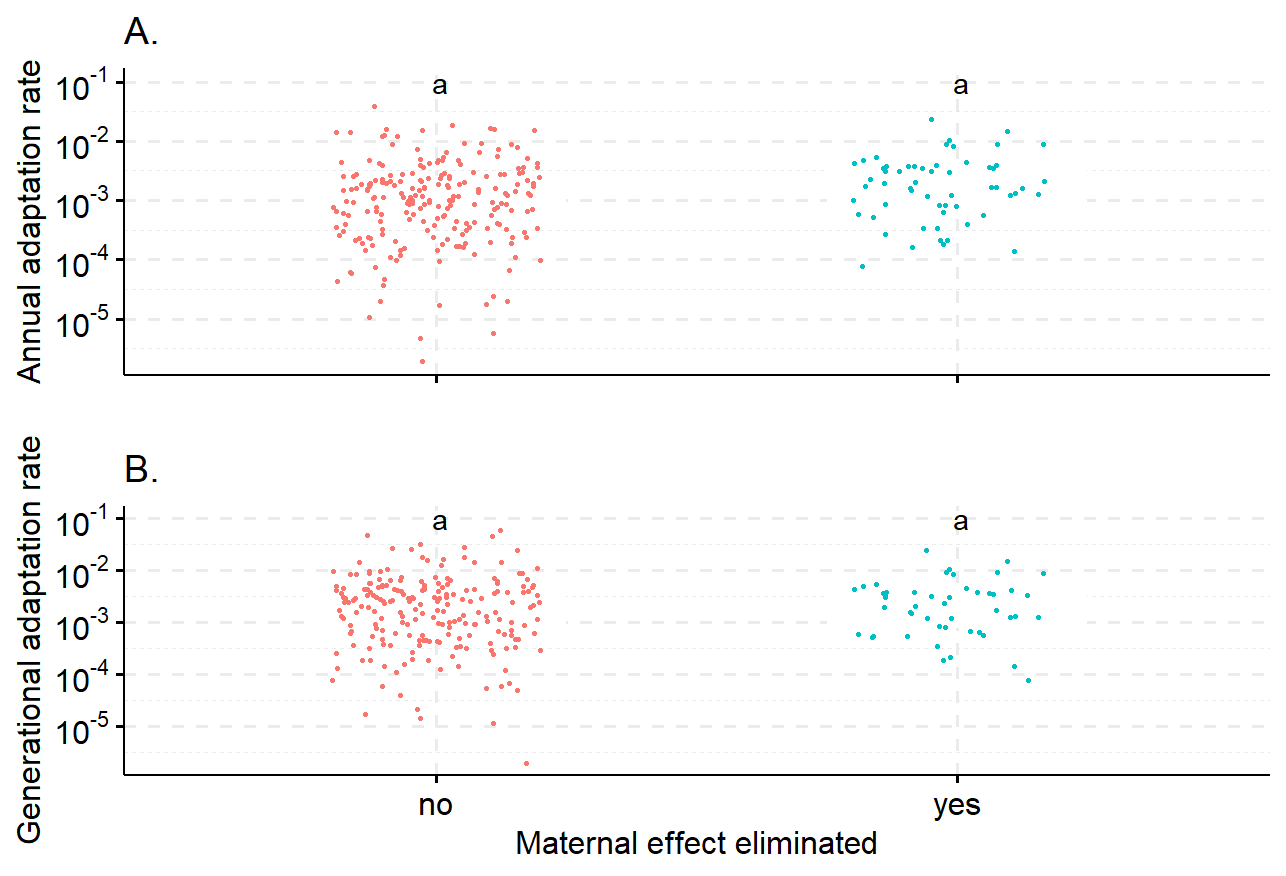

**Fig. S7 Annual (A) and generational (B) adaptation rates of plant traits with different significant levels in the trait variation between the native and invasive plants from the original studies.** ‘Yes’ and ‘No’ indicate those comparisons that were reported as either statistically significant or not (p < 0.05), respectively, in the original study. ‘Unknown’ indicates that the significant levels were not available in the original studies (e.g. the comparisons of treatments that we selected in our database were not shown as a comparison in the original papers). Different lowercase letters indicate that the means of plant annual or generational adaptation rates are significantly different between life history categories (as determined by Tukey post-hoc tests). When significant differences were detected, R^2^ indicates the explained variation, as estimated by Generalized Linear Model (GLM) with Weibull distribution.

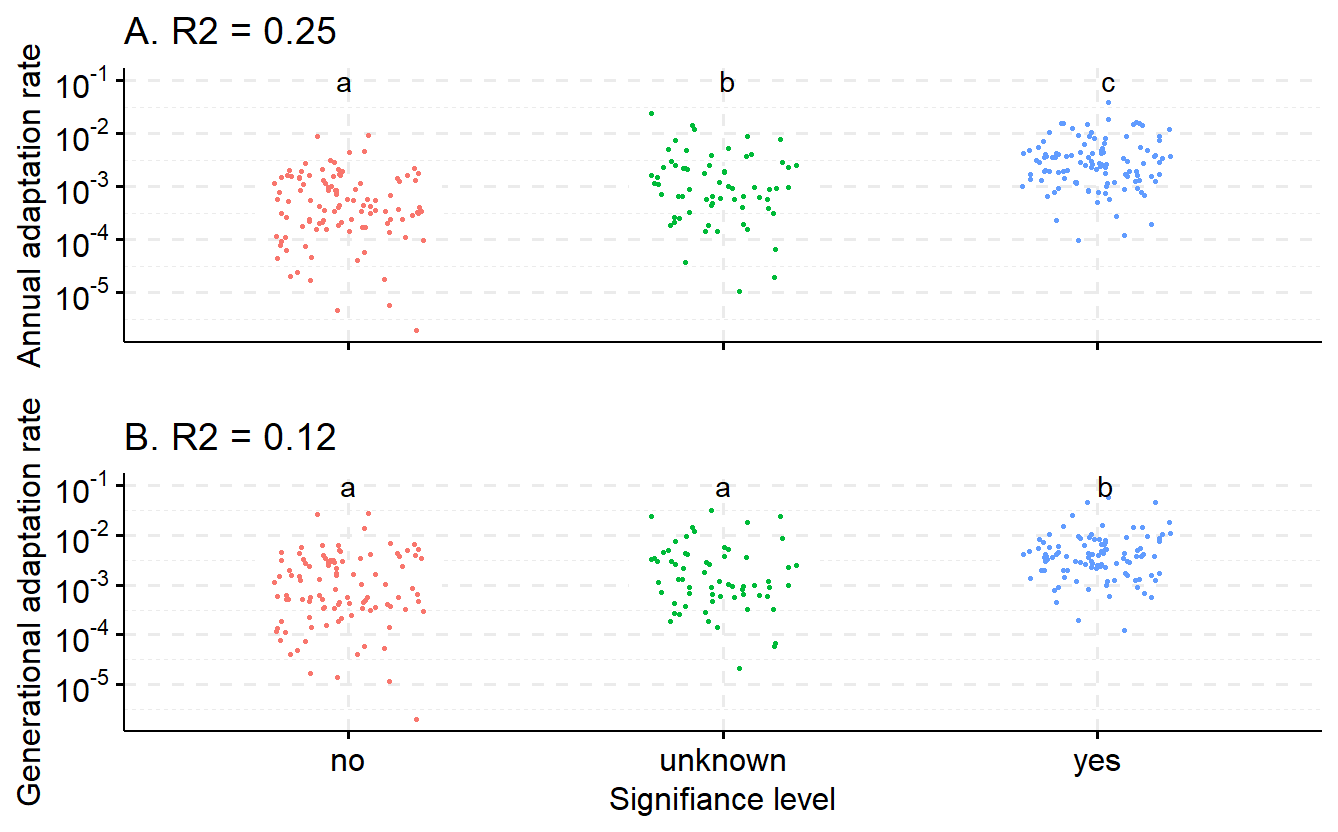

**Fig. S8 Annual (A, B, C) and generational (D, E, F) adaptation rates of plant traits with different significant levels in the trait variation between the native and invasive plants from their original studies split by life history categories** (A & D. generation time less than 1 year, B & E. generation time between 1 and 2 years, C & F. generation time more than 2 years). ‘Yes’ and ‘No’ indicate those native and invasive plant comparisons that were reported as either statistically significant or not (p < 0.05), respectively, in the original study. ‘Unknown’ indicates that the significant levels were not available in the original studies (e.g. the comparisons of treatments that we selected in our database were not shown as a comparison in the original papers). Different lowercase letters indicate that the means of plant annual or generational adaptation rates are significantly different between life history categories (as determined by Tukey post-hoc tests). When significant differences were detected, R^2^ indicates the explained variation, as estimated by Generalized Linear Model (GLM) with Weibull distribution.

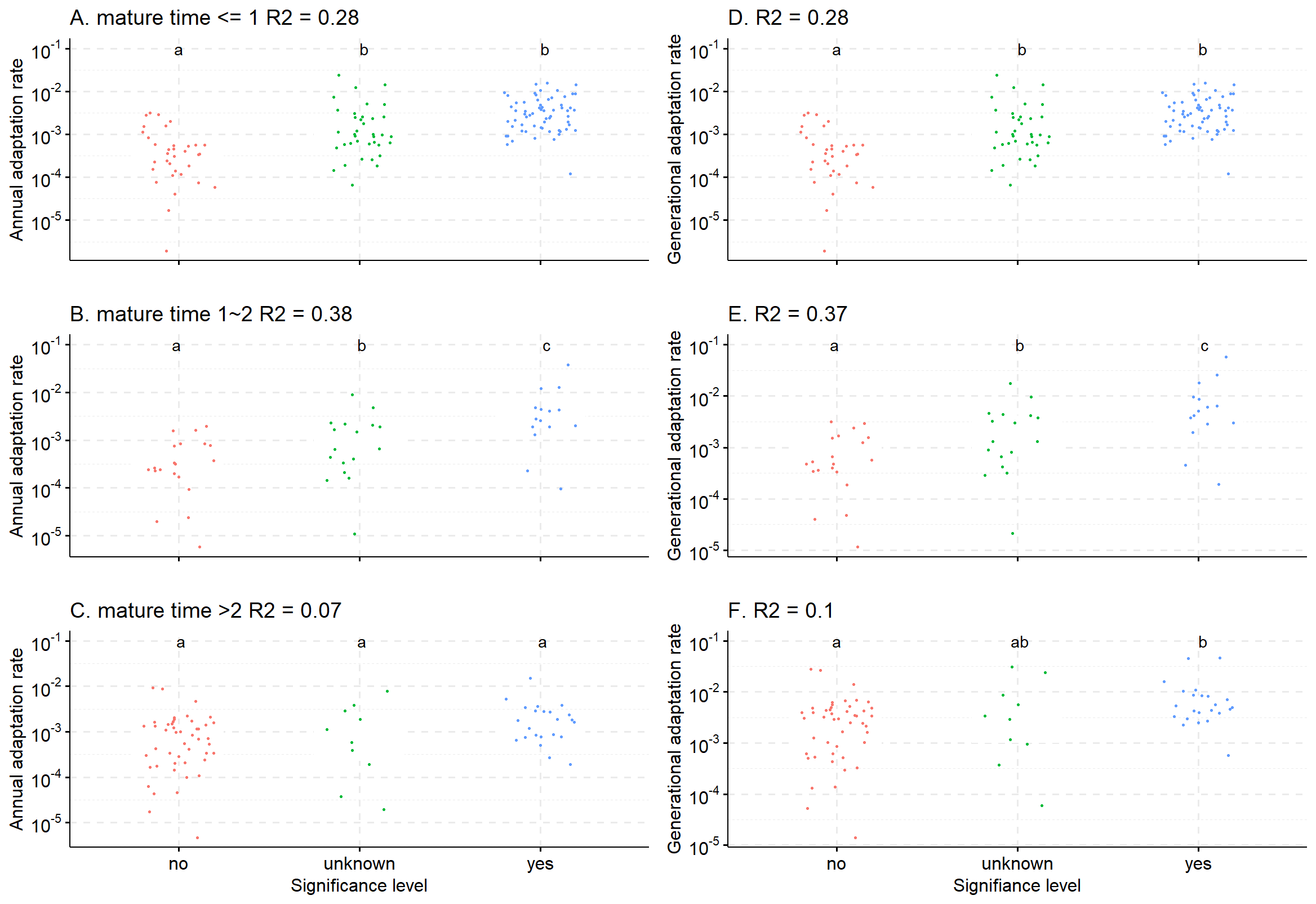

**Results S1 Patterns of annual and generational adaptation rates between growth form, phylogenetic group and trait type split by different life history categories (including Fig. S9 - S11)**

The following figures contain the information shown in the figures (Fig. 4 – 6) in the main text categorised by different life history categories. This allows the assessment of the effect of life history on the variation in adaptation rates caused by growth form (Fig S9), phylogeny (Fig S10) and trait type (Fig S10).

**Fig. S9 Patterns of annual (A, B, C) and generational (D, E, F) adaptation rates of plant traits among different growth forms split by life history categories** (A & D. generation time less than 1 year, B & E. generation time between 1 and 2 years, C & F. generation time more than 2 years). Different lowercase letters indicate that the means of plant annual or generational adaptation rates are significantly different between life history categories (as determined by Tukey post-hoc tests). When significant differences were detected, R^2^ indicates the explained variation, as estimated by Generalized Linear Model (GLM) with Weibull distribution.

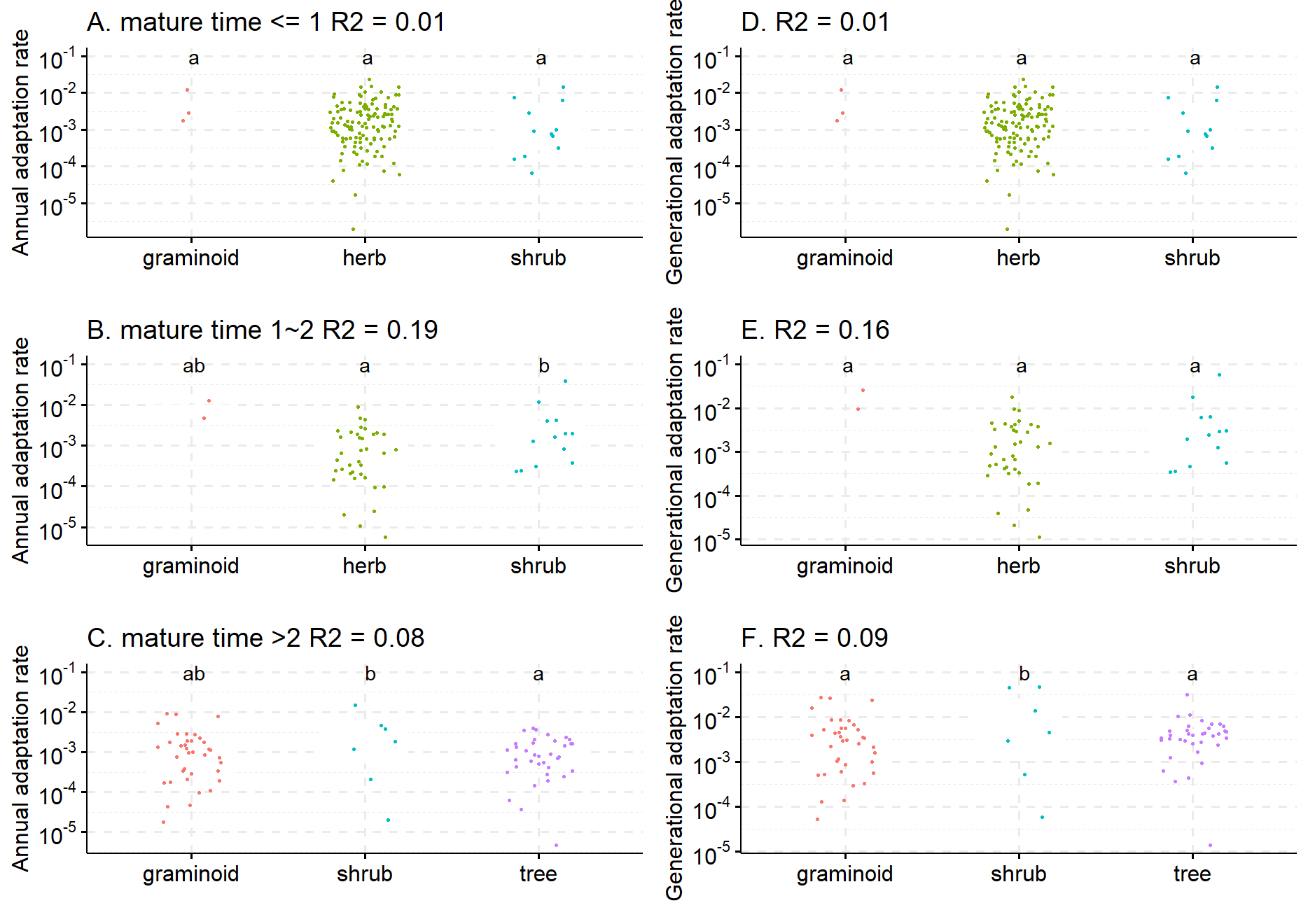

**Fig. S10 Patterns of annual (A, B, C) and generational (D, E, F) adaptation rates of plant traits among different phylogenetic groups split by life history categories** (A & D. generation time less than 1 year, B & E. generation time between 1 and 2 years, C & F. generation time more than 2 years). Different lowercase letters indicate that the means of plant annual or generational adaptation rates are significantly different between life history categories (as determined by Tukey post-hoc tests). When significant differences were detected, R^2^ indicates the explained variation, as estimated by Generalized Linear Model (GLM) with Weibull distribution.
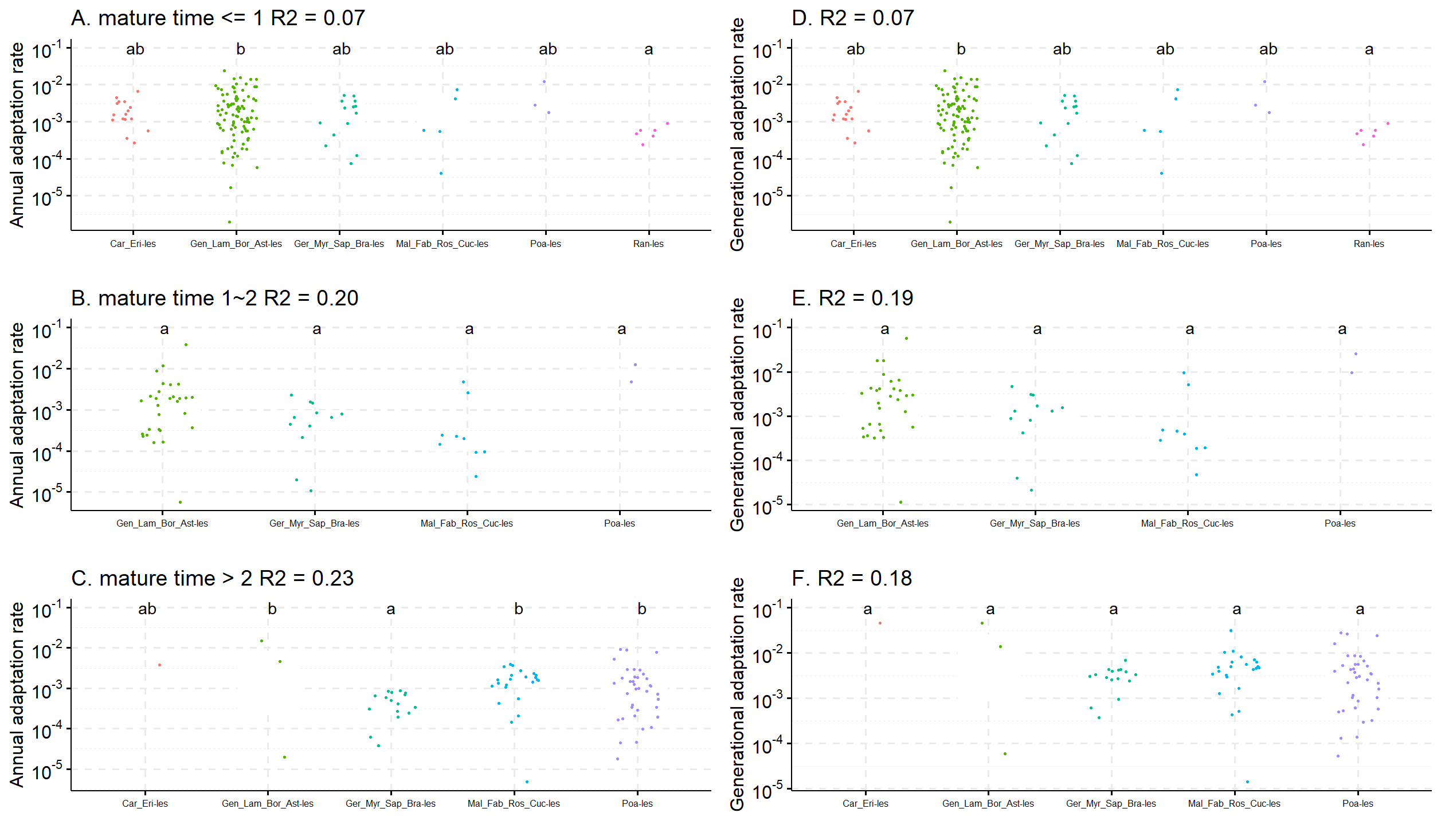

**Fig. S11 Patterns of annual (A, B, C) and generational (D, E, F) adaptation rates of plant traits among different trait groups split by life history categories** (A & D. generation time less than 1 year, B & E. generation time between 1 and 2 years, C & F. generation time more than 2 years). Different lowercase letters indicate that the means of plant annual or generational adaptation rates are significantly different between life history categories (as determined by Tukey post-hoc tests). When significant differences were detected, R^2^ indicates the explained variation, as estimated by Generalized Linear Model (GLM) with Weibull distribution.

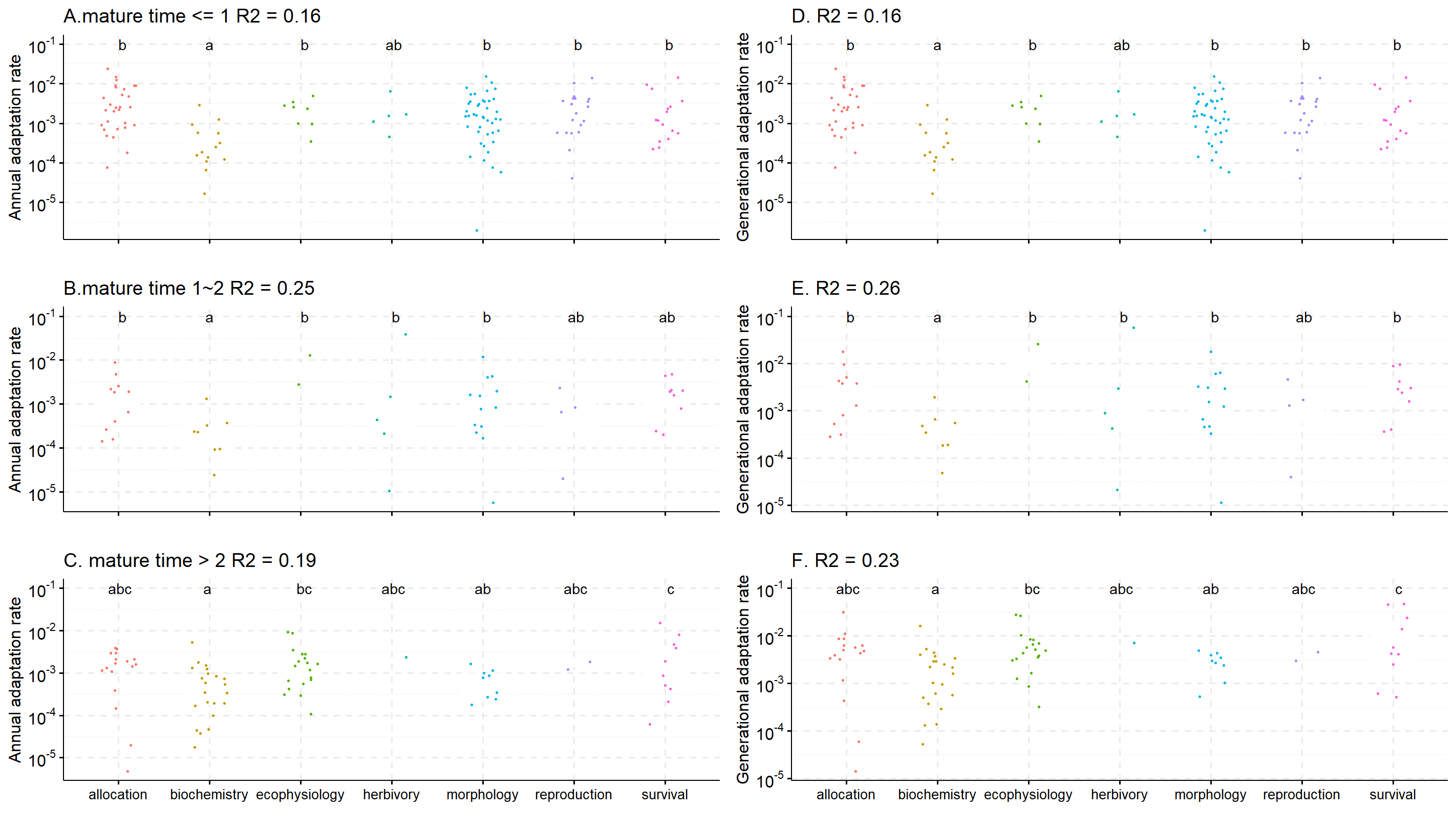
